# MONFIT: Multi-omics factorization-based integration of time-series data sheds light on Parkinson’s disease

**DOI:** 10.1101/2024.06.03.597147

**Authors:** Katarina Mihajlović, Noël Malod-Dognin, Corrado Ameli, Alexander Skupin, Nataša Pržulj

**Affiliations:** Barcelona Supercomputing Center (BSC), 08034 Barcelona, Spain; The Integrative Cell Signalling Group, Luxembourg Centre for Systems Biomedicine (LCSB), University of Luxembourg, Esch-sur-Alzette, Luxembourg; Department of Physics and Material Science, Luxembourg, Luxembourg; University of California San Diego, La Jolla, CA 92093, USA; Department of Computer Science, University College London, WC1E 6BT London, United Kingdom; ICREA, Pg. Lluís Companys 23, 08010 Barcelona, Spain

## Abstract

Parkinson’s disease (PD) is a severe and complex multifactorial neurodegenerative disease with still elusive pathophysiology preventing the development of curative treatments. Molecular deep phenotyping by longitudinal multi-omics is a promising approach to identify mechanisms of PD aetiology and its progression. However, the heterogeneous data require new analysis frameworks to understand disease progression across biological entities and processes. Here, we present MONFIT, a holistic analysis pipeline that integrates and mines time-series single-cell RNA-sequencing data with bulk proteomics and metabolomics data by non-negative matrix tri-factorization, enabling prior knowledge incorporation from molecular networks. First, MONIFT integrates time-point-specific data and then holistically mines the integrated data across time points. By applying MONFIT to longitudinal multi-omics data of differentiation of PD and control patient-derived induced pluripotent stem cells into dopaminergic neurons, we identify novel PD-associated genes, emphasize molecular pathways that play important roles in PD pathology, and suggest new intervention opportunities using drug-repurposing. MONFIT is fully adaptable to other multi-omics data sets.

## 1 Introduction

Parkinson’s disease (PD), the second most prevalent neurodegenerative disorder, is a severe and complex multifactorial disease that affects about 2-3% of the population over the age of 65 [72]. The aetiology of PD is largely unknown, with different genetic mutations (e.g., *PINK1*, *SNCA*, *LRRK2* and *PARK2*) causing approximately 5-10% of all cases, while the remaining cases are considered idiopathic. Although PD is characterized by genetic and clinical heterogeneity, a typical hallmark of PD is the loss of midbrain dopaminergic (mDA) neurons in the substantia nigra of the mid-brain, associated with the accumulation of misfolded *α*-synuclein proteins called Lewy bodies [6].

In our recent study [65], we investigated mDA neurons carrying a *PINK1* mutation, which were differentiated from pluripotent stem cells (iPSCs) and found that PD pathophysiology is characterized by a network-wide systems response of a core set of genes that participate in many interconnected pathways, including, ubiquitination, RNA processing, cellular response to stress, protein metabolism and lysosomal proteins. To get a more comprehensive understanding of the disease mechanisms and to disentangle the complex network response, we need to study PD at a deeper molecular depth by complementary multi-omics analyses. Such multi-omics approaches come with intrinsic challenges to understand their interrelationships of how the regulatory changes captured by the transcriptomics are established on the proteome level and further manifested on the metabolome level.

Hence, there is a need for methods that collectively integrate and mine such multi-omics data to get a holistic view that encompasses the information across the various omics layers and unravel the interactions between disease-associated biomolecules and their functions. Many multi-omics integration tools have been developed that fuse and analyze omics data across different modalities to answer various biological questions related to disease and their mechanisms [79, 37]. Earlier integration tools have focused on fusing bulk multi-omics data [9, 79], but the explosion of the single-cell (SC) data has given rise to algorithms that do so at the SC level, most prominently for SC transcriptomics (i.e., scRNA-seq) datasets [56, 37]. To guide the integration procedure to biologically meaningful results and complement the information of experimental omics measurements to discover new biology, multiple integration methods also incorporate molecular interaction networks, such as protein-protein interaction (PPI) or coexpression (COEX) networks available from various bioinformatics databases, as prior knowledge (e.g. [12, 53, 47]). A powerful and popular approach for data fusion is non-negative matrix factorization (NMF) method [50, 25], which decomposes multiple omics modalities represented as input matrices into products of two matrix factors that could be interpreted as embeddings [25]. NMF-based and its more recent extension of non-negative matrix tri-factorization (NMTF) allow for an intuitive and easy interpretation of the resulting matrix factors [64, 2, 39, 34, 23] and have been used to study molecular interaction networks and uncover novel biological mechanisms [73, 58, 86]. However, no method is designed to integrate multi-omics bulk and scRNA-seq data with molecular interaction networks and fully exploit such integrated data in a time-series setting for downstream tasks of uncovering novel disease-associated genes, highlighting important disease pathways, and proposing new intervention strategies.

To overcome these limitations, we here adapt our integration and downstream mining framework from Mihajlovic *et al.* [61] to propose MONFIT, a **M**ulti-**O**mics **N**on-negative matrix tri-**F**actorization **I**ntegration of **T**ime-series data. First, MONFIT uses NMTF to integrate multi-omics time-point-specific data of scRNA-seq, bulk proteomics, and bulk metabolomics measurements of disease and control samples from a time-series experiment with prior knowledge from molecular interaction networks in terms of PPI, metabolic-interaction (MI), genetic interaction (GI), and COEX, producing gene embeddings. Then, MONFIT holistically mines these embeddings across all time points to identify new disease-associated genes, by applying a new downstream method based on the concept introduced in Mihajlovic *et al.* [61], which states that those genes whose position changes the most between disease and control samples across all time points are disease-related.

We use MONFIT to study the differentiation dynamics of a patient-derived PD cell line harbouring a *PINK1* mutation in comparison to a matching control by integrating scRNA-seq, bulk proteomics, and bulk metabolomics data with molecular interaction networks, predicting 163 genes that are globally related to PD in the literature and are specific to the *PINK1* mutation causing PD. We compare our gene predictions with differentially expressed genes (DEGs), and differentially abundant proteins (DAPs) obtained from the original study of the data [8] and discover a common set of prioritized genes in all three analyses, which indicates their importance in the pathology and development of PD. We further demonstrate that MONFIT goes beyond standard differential analysis approaches of single-omics data by predicting PD-associated genes that would otherwise elude discovery. By performing enrichment analysis, we show that our 163 predictions participate in pathways pertinent to PD, highlighting multiple pathways that could serve as new intervention points in PD. Moreover, we manually validate the top 30 predictions in the literature, finding five genes (*CENPF*, *CRABP1*, *TOP2A*, *TMSB10* and *NASP*) that had no known PD association and provide compelling evidence elucidating their role in PD. Finally, we perform an enrichment analysis in drug-target interactions from DrugBank [88] to propose future therapeutic strategies based on drug repurposing (e.g., Artenimol).

## 2 MATERIALS AND METHODS

### 2.1 Multi-omics datasets of Parkinson’s disease iPSC model

We collect the longitudinal data set containing scRNA-seq, bulk metabolomics, and bulk proteomics of iPSC-derived time-series cell lines of a *PINK1* mutation causing PD and a corresponding control at five time points; day 8, 18, 25, 32 and 37 (denoted by D8, D18, D25, D32 and D37, respectively) [8]. From this data set, we obtain a multi-omics characterization of ten cell conditions, where each condition is defined by a cell type (*PINK1* mutation or control) at a specific time point.

From Bernini *et al.* [8], we collect scRNA-seq data, which has undergone quality control. We model the expression data of each cell condition by a matrix *E* in which rows represent genes, columns represent cells, and an entry *E_ij_* is the raw read count of a gene *i* in a cell *j*. Next, we normalize each count matrix by applying the shifted logarithm for stabilizing the variance for the subsequent dimensionality reduction analysis. Finally, we filter the condition-specific transcriptomics data, keeping protein-coding genes with at least one PPI in BioGRID, as PPIs are the most direct evidence that two genes interact (see Supplementary Table 2 for the size of each *E* matrix). Emerging evidence indicates that disrupted energy metabolism plays a key role in PD [1, 11] and that analyzing metabolic interactions facilitates the discovery of novel PD-associated genes [61]. Thus, we obtain metabolomics data that include metabolite abundance of 70 metabolites from liquid and 58 metabolites from gas chromatography-mass spectrometry across all cell conditions from Bernini *et al.* [8]. First, we scale each metabolomics dataset with the mean of non-zero values to reduce the impact of extreme values and bring the datasets to a comparable scale. Next, for each metabolite in a dataset and cell condition, we average the abundance across replicates [32] in the following way: i) if no replicate is 0, we compute the mean across all replicates, ii) if *≤* half of the replicates are non zero, we compute the mean of the non-zero replicates, or iii) otherwise, we set the average abundance to zero. This method helps to mitigate the impact of missing values while preserving as much information as possible from replicates with non-zero values. Finally, we merge the two datasets to obtain one metabolite abundance matrix. As both methodologies measure some of the same metabolites, we choose the abundance of such metabolites from one measurement type based on the highest mean value across all cell conditions. This strategy ensures that the measurement of the metabolite abundance obtained by the chromatography-mass spectrometry method, which is more sensitive for quantifying that specific metabolite, is used [77]. This results in the metabolomics matrix containing metabolite abundances of 96 metabolites.

To gain novel molecular insights into pathological mechanisms underlying PD and identify new PD biomarkers, promising findings were uncovered by analyzing bulk proteomics data [70, 43]. Thus, we obtain a bulk proteomics dataset that contains the protein abundance measured by liquid chromatography-mass spectrometry for each cell condition from Bernini *et al.* [8]. Protein IDs in proteomics data are mapped to gene symbols using SynGO [46]. Outlier replicates are filtered out by using hierarchical clustering, where only 2 out of 21 replicates are excluded. To ameliorate the technical artifacts of the measuring technique, we impute the missing values by using FactoMineR package [49]. Then, we average the protein abundance of each protein by applying the same approach used for metabolites (described above). In addition, we remove duplicate proteins’ lower measured protein abundance and log transform the resulting proteomics matrix to minimize the impact of outliers [14]. This results in the proteomics matrix containing the protein abundance of 7380 proteins.

### 2.2 Constructing molecular networks

We collect four general molecular interaction networks (PPI, MI, COEX and GI) for *Homo sapiens* (human) that represent prior knowledge (detailed below). To incorporate the data with prior knowledge and produce data-driven molecular networks, we create (i) condition-specific PPI networks that account for measured protein abundances, (ii) condition-specific gene-centric MI networks that account for metabolite abundances, (iii) condition-specific GI networks, and (iv) condition-specific COEX networks, as follows.

We create a PPI network by collecting all physical interactions between proteins from BioGRID database version 4.4.218 [68], captured by at least one of the following experiments: Two-hybrid, Affinity Capture-Luminescence, Affinity Capture-MS, Affinity Capture-RNA, Affinity Capture-Western. For each cell condition, we generate a condition-specific subgraph of the PPI network obtained from BioGRID and induced by the genes expressed in that cell condition (as measured by scRNA-seq). To incorporate the proteomics data in the condition-specific PPI networks, we set the weight of an edge between two genes as a product of their corresponding protein abundances measured for each cell condition. The intuition behind this approach is that the expected interaction between two genes depends on the abundance of their protein products, so the more abundant the two protein products are, the more likely the interaction between them occurs. For genes that are expressed and can interact in a condition-specific PPI but whose protein abundance is not measured by the proteomics data, we set the edge weight between them as the smallest product between the proteins measured in a condition. This approach assigns higher weights to the interactions between genes whose protein product abundances have been detected and quantified while minimizing and preserving the weights of the remaining interactions because the unmeasured protein abundances may result from technical noise rather than the absence of such proteins.

For each cell condition, we construct a gene-centric MI network so that an edge connects two enzyme-coding genes if any product metabolite of the first enzyme is the substrate metabolite of the second enzyme [73]. Because enzymes can be defined by more than one gene, we consider that all the genes defining one enzyme interact with all the genes defining the other enzyme (all-to-all interactions). Each condition-specific MI network is composed of genes that form condition-specific PPI networks. We collect information that relates enzymes, genes and metabolites from KEGG 105.0 release [42]. In contrast to the proteomics data, where the protein abundance is tied to the nodes (i.e., genes) of the network, the metabolite abundance can be directly seen as an edge weight (i.e., interaction strength) between two genes. Thus, to incorporate the information in the condition-specific metabolomics data with the MI networks, we set the weight of an edge between two interacting genes as the sum of all metabolites that participate in the enzyme-enzyme interactions related to these genes. Due to the targeted measurement of metabolomics data, only a small subset of metabolites that facilitate the MIs have been quantified. To account for the lack of measurements, for metabolites whose abundance is not quantified by the metabolomics data, we set their value as the median metabolite abundance value of the metabolites measured in a condition. To make the condition-specific GI network, we collect all genetic interactions reported in Bi-oGRID 4.4.128 [68] between genes constituting condition-specific PPI networks. Finally, we create condition-specific COEX networks between genes forming the condition-specific PPI networks by collecting gene interactions from CoexpressDB v.8 [66], whose z-score is *≥* 3, thereby capturing the top 1% of the strongest interactions.

In the following, we represent molecular networks by their adjacency matrices, *A*_*i*∈{1,…,4}_, where each *A_i_* is a symmetric matrix. Each of our condition-specific GI and COEX networks are unweighted networks, so their adjacency matrices are binary matrices *A*_1_ and *A*_2_, respectively. An entry value *A_i_*[*v, w*] for *i* ∈ {1, 2} of one between two interacting genes *v* and *w* indicates that genes *v* and *w* interact, and zero otherwise. On the other hand, our condition-specific PPI and MI networks are weighted networks represented by adjacency matrices *A*_3_ and *A*_4_, respectively, where a positive entry value *A*_3_,_4_[*v*][*w*] between two interacting genes *v* and *w* is the weight of the corresponding interaction edge, and zero indicates that two genes do not interact.

### 2.3 Data integration

After characterizing each cell condition with five layers of multi-omics data (data-driven GI, COEX, PPI and MI networks, and scRNA-seq data; left panel of Figure 1A), we perform data integration by adapting and applying our NetSC-NMTF integration framework from Mihajlovic *et al.* [61] (right panel of Figure 1A). For each cell condition, NetSC-NMTF takes as input adjacency matrices, *A_i∈{_*_1_*_,…,_*_4}_, representing molecular interaction networks, and an SC expression matrix, *E*, simultaneously decomposing them into products of three matrix factors. Thus, 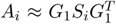 for all *i* and 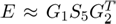, where *G*_1_ *∈* ℝ^*n*×*k*_1_^, *G*_2_ *∈* ℝ^*m*×*k*_2_^, *S_i∈{_*_1_*_,…,_*_4_*_}_ ∈* ℝ^*k*_1_×*k*_1_^ and *S*_5_ ∈ ℝ^*k*_1_×*k*_2_^, with *n* being the number of genes, *m* the number of SCs, and *k*_1_ and *k*_2_ the dimensions of the latent embedding spaces *G*_1_ and *G*_2_, respectively. As *G*_1_ and *G*_2_ can be interpreted as cluster indicator matrices of genes and SCs, respectively, we choose the number of dimensions *k*_1_ and *k*_2_ by using the rule of thumb, 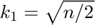 and 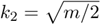, which is a heuristic used to determine a reasonable number of clusters given the number of genes or SCs that need to be clustered [45] (see Supplementary Table 3 for values of *k*_1_ and *k*_2_ of each cell condition). In addition, we demonstrate that our adapted NetSC-NMTF integration framework is robust to the choice of *k*_1_ and *k*_2_ parameters (Supplementary Section 1). As *G*_1_ is shared across all matrix decompositions, it facilitates information flow, allowing the integration procedure to learn from all data. *S_i∈{_*_1_*_,…,_*_4}_ matrices are the compressed representations of the molecular networks, and *S*_5_ is the compressed representation of the SC expression matrix, *E*.

**Figure 1:**
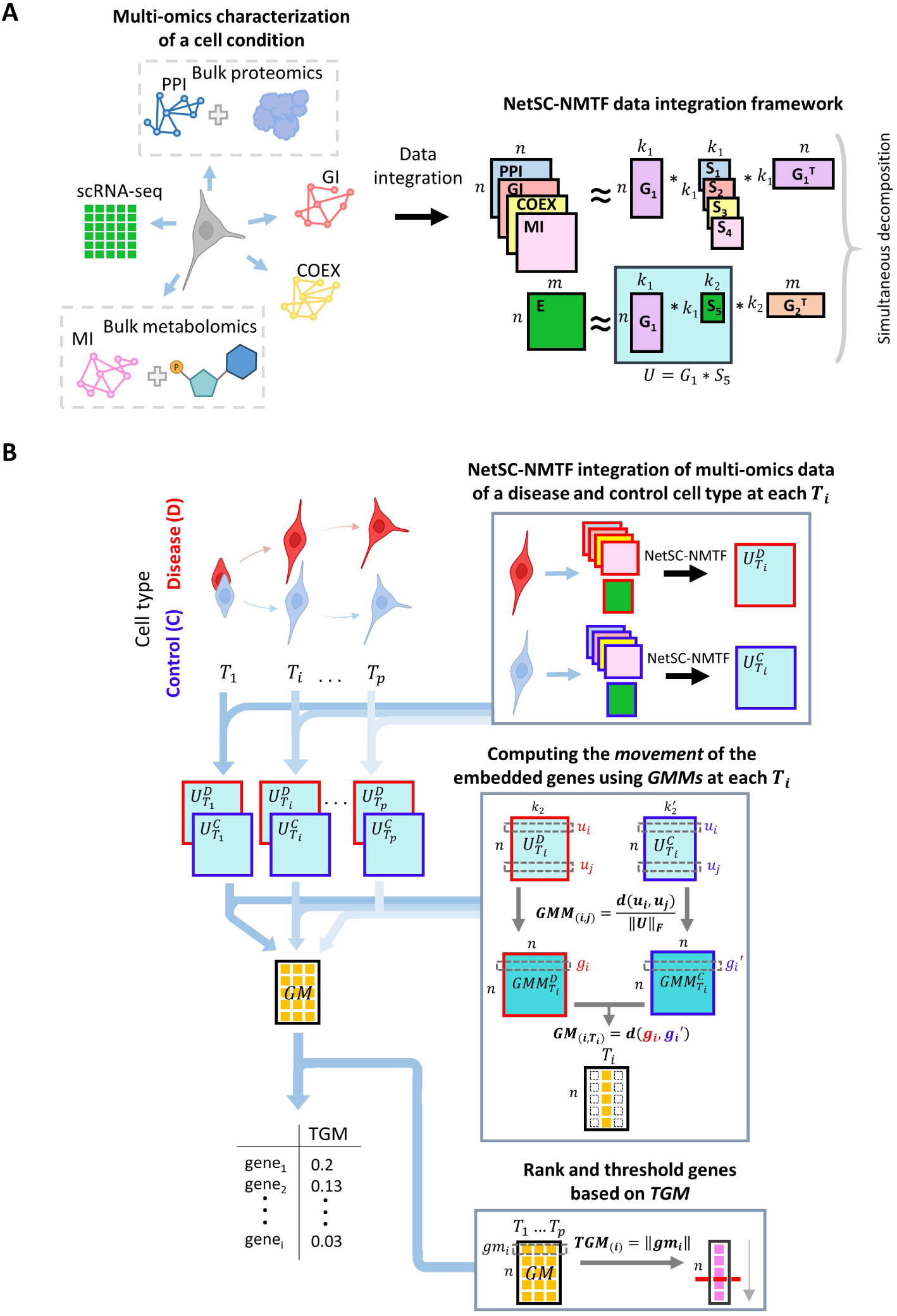
MONFIT pipeline. **(A)** A cell condition is characterized by different multi-omics data that include molecular networks (PPI, MI, GI, and COEX), bulk proteomics and metabolomics, and scRNA-seq data (E). We filter each condition-specific input data matrix, keeping the genes that are in the overlap of protein-coding genes from the PPI network and genes expressed in a cell condition. Furthermore, bulk proteomics (or metabolomics) and a prior knowledge PPI (or MI) network are combined to create data-driven PPI (or MI) networks. We integrate the data by adapting and applying an NMTF-based model called NetSC-NMTF [61] to obtain the gene embeddings (*G*_1_), which we map to the space of SCs with the transformation matrix *S*_5_, obtaining gene embeddings captured by matrix *U*. **(B)** To compare multi-omics time-series data between disease and control, we propose MONFIT. First, MONFIT individually integrates the multi-omics data of a disease and control cell condition at one time point using the integration approach from **(A)**, obtaining paired disease and control gene embeddings (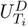 and 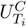). Second, MONFIT computes the relative position of each gene against all the other genes for a given disease condition by using the norm-scaled Euclidean distances between their embedding vectors from 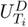, resulting in a 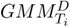. MONFIT performs the same step for the control embeddings 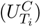, obtaining 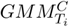. Then, it compares the relative position of each gene in 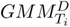, with its relative position in 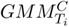 to determine the “movement” of a gene between a disease and control cell condition at one time point. By computing this “gene movement” of all genes across all time points, MONFIT obtains the *GM* matrix whose dimensions define an embedding space that captures “gene movement” across time. Finally, to prioritize genes, MONFIT computes each gene’s “total gene movement” (*TGM*) by computing the length of the vector of “gene movements” across time, ranking the genes according to the highest *TGM*. Finally, MONFIT obtains disease-related predictions by thresholding the ranked genes, focusing on the top-moving ones.

As solving NMTF is an NP-hard continuous optimisation problem [73], we obtain all matrix factors by applying a heuristic fixed-point solver based on multiplicative update rules (MURs) [24] (see Mihajlovic *et al.* [61] for details and derivation of MURs) by minimising:

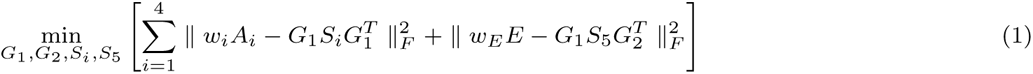

where *G*_1_*, G*_2_ *≥* 0, ‖ ‖*_F_* denotes the Frobenius norm, and *w_i_* and *w_E_* are weighting factors that determine how much each input matrix contributes to the integration procedure. Starting from an initial solution, the solver iteratively employs MURs to converge towards a locally optimal solution. To initialize the matrix factors, we apply singular value decomposition (SVD) on the original input matrices because it reduces the number of iterations required for convergence and leads to a deterministic solution [74]. We take the absolute values of the entries of the computed SVD matrices to comply with the non-negativity constraint of NMTF. The iterative minimization process stops when the objective function converges, which we evaluate every ten iterations with: 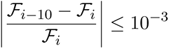, where ℱ is the value of the objective function at the current iteration and ℱ*_i−_*_10_ is the value computed at iteration *i −* 10.

### 2.4 Using matrix factors to define gene embeddings

According to the embedding interpretation of NMTF, we interpret the rows of *G*_1_ matrix as *k*_1_-dimensional embedding vectors of genes (i.e., gene embeddings), and the rows of *G*_2_ matrix as *k*_2_-dimensional embedding vectors of SCs. In addition, *S*_5_ can be interpreted as a transformation matrix, coupling the underlying biology of a cell condition captured by the multi-omics measurements with its SC phenotype. Therefore, to emphasize the connection between the multi-omics data and the resulting SC phenotype, we project the gene embeddings from *G*_1_ to the space spanned by *G*_2_ by using the transformation matrix *S*_5_; we compute these projected gene embeddings, *U*, as *U* = *G*_1_*S*_5_.

In Sections “Finding the optimal combination of data for integration” and “The optimal combination of data for integration”, we study the functional organization of gene embeddings in *G*_1_ to determine the best combination of input matrices for integration. The intuition behind this approach is that gene embeddings in *G*_1_ matrices are functionally organized so that genes close together in the gene embedding space participate in the same biological mechanisms. Therefore, this allows for identifying which combination of input data produces gene embeddings whose organization best captures the disease-related biological mechanisms of individual cell conditions. In contrast, *U* gene embeddings are better at capturing the differences between the cell conditions because projecting gene embeddings into the space of SCs highlights the biological signal caused by different SC phenotypes. Therefore, we compare *U* gene embeddings between PD and control cell conditions to obtain new PD-associated gene predictions.

### 2.5 Finding the optimal combination of data for integration

A general challenge in integrating multi-omics data is the optimal incorporation of heterogeneous data for biologically interpretable embeddings. Following the idea that genes with similar embeddings participate in the same molecular mechanisms [52], we first determine which combination of the five input matrices produces gene embeddings that capture best the molecular mechanisms related to PD. In this respect, we refer to gene embeddings as the rows of *G*_1_ matrix obtained by applying the first step of the MONFIT pipeline. For each cell condition, we define the combination of data by a 5-tuple of weights (*w_P P I_*, *w_COEX_*, *w_GI_*, *w_MI_*, *w_E_*) that represent the weighting factors by which the corresponding input matrices (PPI, COEX, GI, MI and E) are multiplied before applying NetSC-NMTF to obtain gene embeddings. We test all combinations *w_P P I,COEX,GI,MI_ ∈* {0, 0.1, 1, 10*}*^4^ and *w_E_ ∈ {*0.1, 1, 10*}* for the SC expression matrix weights resulting in 768 combinations. All those ranges allow for not having these datasets except SC expression, which is the simplest data that we analyze.

First, we cluster the gene embeddings of each combination and cell condition by applying 10 runs of k-means clustering (where the number of clusters is the same as the dimension of the embedding space). Second, we perform enrichment analysis of these clusters in PD-related mechanisms based on pathways from the PD map [29] by applying a hypergeometric test (Supplementary Section 2) and measuring the percentage of enriched clusters and of enriched genes. We exclude *Scrapbook* and *Parkinson’s UK Gene Ontology genes* from the terms collected from the PD map because they do not represent biological pathways. Third, for each weight combination and cell condition, we compute the mean of the percentage of enriched clusters and of enriched genes across the k-means runs. Next, for each weight combination, we compute the median of each enrichment measurement across all cell conditions (by relying on the median, we do not assume the shape of the distribution of the enrichment values across the cell conditions). We assign a rank to each weight combination according to the median percentage of enriched clusters and enriched genes, so that the higher the enrichment is, the lower the rank. Finally, we determine the overall ranking by computing the average rank score between the two rankings and selecting the top-ranked combination (i.e., with the lowest rank) of input data to use for integration.

To further assess the biological relevance of the condition-specific gene embeddings obtained with the most optimal combination of input data, we perform enrichment analysis on the gene clusters obtained with k-means in biological annotations of GO biological processes (GO-BP) [4], KEGG [42] and Reactome pathways [41], using a hypergeometric test (Supplementary Section 2).

### 2.6 MONFIT pipeline

To study the multi-omics time-series data of a PD cell line compared to control to predict new genes that potentially cause or drive the progression of PD, we propose MONFIT (pipeline illustrated in Figure 1B). MONFIT is a 3-step approach that builds upon the method presented in Mihajlovic *et al.* [61], which is a method that uncovered valuable biological insights in the context of PD by integrating and mining multi-omics data from a time-series experiment. In summary, the method from Mihajlovic *et al*. [61] predicts novel PD-associated genes by applying the NetSC-NMTF framework to obtain “gene embeddings” of time point-specific data. Then, the embeddings are mined using the following 2-step downstream method. In the first step, for each time point, the method uncovers gene predictions that are statistically significantly associated with known PD genes, i.e., genes that appear in the clusters that are significantly enriched in known PD genes. In the second step, the method defines the final set of gene predictions as genes that are statistically significantly associated with known PD genes at all time points, which are obtained by intersecting all sets of time-point-specific predictions from the first step. Finally, the final set of gene predictions is prioritized by computing the average changes in gene embeddings between control and disease conditions across all time points, ranking the ones with the highest average change on top.

MONFIT builds upon these methods, improving them as follows. In the first step, MONFIT applies the adapted NetSC-NMTF framework (Section “Data integration”), which allows for the integration of multi-omics data from each condition (disease or control cells at each time point) separately, producing condition-specific gene embeddings. The adapted NetSC-NMTF is different from NetSC-NMTF originally introduced in Mihajlovic *et al.* [61] because it allows for: i) integrating bulk proteomics and metabolomics data through the data-driven PPI and MI networks and ii) regulating the relative contribution of each omics data on the resulting gene embeddings through weighting factors. Incorporating bulk proteomics and metabolomics data during integration is beneficial as it leads to a reduced set of PD predictions that are more indicative of initial cellular response to the *PINK1* mutation and potential PD causality (see Supplementary Section 4 for a comparison between the predictions obtained by including and excluding bulk omics data during integration). By enabling the variability of the weighting factors, we allow the method to choose which omics data contributes to the organization of genes that best capture disease biology. In the second step, for each time point, MONFIT measures how the embeddings of genes change between the corresponding PD and control conditions. We term these changes as the genes’ “movement”. In contrast, in Mihajlovic *et al.* [61], the downstream mining method of the gene embeddings first focuses on genes that cluster with known PD genes across time points, and prioritizes them based on their “movement”. The advantage of MONFIT’s second step is that it does not bias the discovery of new PD-associated genes to only those genes that are close in the embedding space to known PD genes, thereby enabling the identification of PD-related genes that might still contribute to disease development, but do not necessarily cluster together with PD genes.

Finally, in the third step, MONFIT collectively mines the gene “movement” across all time points by computing an overall “movement” that is more robust to outliers (genes that exhibit only one very large change in one time point, rather than changes at all time points) than the mean of gene “movements” across time points presented in Mihajlovic *et al.* [61]. This uncovers genes that potentially cause or drive the progression of PD across all time points and are more directly related to the *PINK1* mutation causing PD (Supplementary Section 5). In the following, we present each step of MONFIT in more detail.

As the first step, MONFIT applies the adapted NetSC-NMTF integration framework on the multi-omics data of a PD cell condition at a time point (Section “Data integration”), *T_i_*, and a control condition at *T_i_*. This results in one set of gene embeddings corresponding to a disease state, 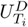, and another set of gene embeddings of control, 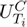. MONFIT repeats this for all *p* time points obtaining a pair of PD and control gene embeddings for each time point (top of Figure 1B). To obtain novel PD gene predictions, MONFIT analyzes these embeddings using the following downstream mining approach.

To understand how PD alters the functional organization of genes compared to a healthy state, we rely on the observation that the gene embeddings of known PD genes (i.e., genes collected from DisGeNet that are associated with Parkinson’s Disease, whose Concept Unique Identifier is C0030567) [71] compared to background exhibit differences between PD and control conditions at matching time points [61]. Computing these differences through a direct comparison between a gene’s embedding vector in 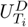 and its embedding vector in 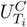 is mathematically incorrect because the coordinates of these two vectors depend on the dimensions that define a basis of an embedding space, which is specific for each cell condition. To overcome this barrier, in the second step, MONFIT applies the methodology introduced in Mihajlovic *et al.* [61] and illustrated in the middle of Figure 1B, where the gene embeddings of each cell condition are transformed into the relative positions of each gene to all other genes, enabling direct comparison between genes of the time-matching disease and control conditions. First, MONFIT converts gene embeddings, matrix *U*, into a symmetric distance matrix called *Gene Mapping Matrix* (*GMM*) that captures the relative positions between the gene embeddings of a cell condition at a particular time point. MONFIT computes a *GMM* with *GMM*_(_*_i,j_*_)_ = *d*(*u_i_, u_j_*)*/*‖*U* ‖*_F_*, where each entry corresponds to the norm-scaled Euclidean distance between the gene embeddings *u_i_* and *u_j_* of two genes *i* and *j* in the matrix *U*, ‖ ‖*_F_* denotes the Frobenius norm and *GMM ∈* ℝ*^n×n^*. Therefore, for a gene *i*, the row *i* of the *GMM*, denoted by *g_i_*, is the relative position of a gene *i* to all the other genes in a cell condition. Then, MONFIT compares each gene between time-matching disease and control *GMMs*, by computing its “gene movement”, which is the change of relative position of a gene between the corresponding disease and control conditions, which we measure as 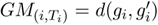, where *d*() denotes Euclidean distance, and *g_i_* and 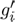 are the relative gene positions in disease and control time-matching *GMMs*, respectively. The “gene movement” is either a positive value and indicates to what extent PD alters the relative position of a gene in a PD embedding space compared to its position in a corresponding control, or zero if there is no such change. By calculating the “movement” of all genes between PD and control cell conditions across all time points, MONFIT obtains a *Gene Movement* matrix (*GM*), where *GM ∈* ℝ*^n×p^*, indicating that each column is an *n*-dimensional vector of “movements” of genes at one time point and each row, *gm_i_*, is a *T*-dimensional vector of “movements” of one gene across all time points; the rows of matrix *GM* can also be viewed as embeddings of genes in an embedding space whose dimensions correspond to the “movements” of genes at individual time points.

To predict novel PD-associated genes that drive the progression of PD during cell development, in the third step, MONFIT identifies which genes moved the most across all time points. For each gene, MONFIT computes the length of its vector of “movements” across time, *gm_i_*, obtaining the “total gene movement” (*TGM*) with *TGM*_(_*_i_*_)_ = ‖*gm_i_*‖. Then, MONFIT ranks the genes according to the highest *TGM* and thresholds them based on the elbow point of the curve of ranked *TGM* values, keeping only the top moving genes (bottom of Figure 1B). MONFIT determines the elbow point using the function KneeLocator as implemented in the package Kneed (v.0.8.5) with a sensitivity parameter of 1, thereby capturing the most moving genes.

## 3 RESULTS AND DISCUSSION

### 3.1 The optimal combination of data for integration

An overarching challenge to integrate multi-omics data lies in effectively incorporating diverse data types to generate biologically interpretable embeddings. To ensure effective data integration of scRNA-seq data, bulk proteomics and metabolomics, and molecular networks, we investigate which combination of the input data results in gene embeddings that best capture the molecular mechanisms associated with PD. Thus, we perform a grid search of the weighting factors *w_P P I_*, *w_COEX_*, *w_GI_*, *w_MI_* and *w_E_* that scale the entries of the input matrices corresponding to PPI, COEX, GI, MI and E, and apply the first step of MONFIT, obtaining gene embeddings for each combination. We determine the best weighting scheme by performing a cluster and enrichment analysis in PD-related pathways from PD map [29] and rank the data combinations according to the highest average rank across the cluster and gene enrichments (Section “Finding the optimal combination of data for integration”).

Overall, we observe a high variability in the percentage of enriched clusters and genes in PD map pathways (Supplementary Figure 1A) across the combinations of weights, indicating the importance of determining the right combination of data. However, by performing a robustness analysis measuring the agreement between the *GMM* matrices of each cell condition across the top 100 ranking combinations of weights, we observe small differences between the *GMMs*, implying that our method is robust to the choice of the weights (Supplementary Section 1). The highest average rank is achieved for the combination of input data with weights of *w_P P I_* = 1, *w_COEX_* = 1, *w_GI_* = 0, *w_MI_* = 10, *w_E_* = 10, which corresponds to 22.86% of clusters with enriched PD map pathways, and 18.07% of genes enriched in PD map pathways (Supplementary Figure 1B). Interestingly, the best weighting factor for the GI network is 0, implying that its information content is not necessary for producing gene embeddings that capture PD biology.

We further assess if the condition-specific gene embeddings obtained with the most optimal combination of weighting factors are globally biologically coherent by performing a cluster and enrichment analysis in GO-BP terms, KEGG (KP), and Reactome (RP) pathways. Across all annotation terms, we find that over 89% of clusters exhibit enrichment in at least one annotation, while for gene enrichments, we observe 34.6%, 44.7%, and 48.1% of genes that are enriched in at least one of their annotations in GO-BPs, KPs, and RPs, respectively (Supplementary Figure 2), indicating the functional relevance of these gene embeddings.

### 3.2 Total gene movement predicts new PD genes

Given the optimal embeddings obtained from the first step of MONFIT, we assess if it is applicable to use “gene movement” to predict new PD-associated genes computed by using the second step of MONFIT. We compare the “gene movements” of known PD genes (i.e., genes obtained from DisGeNet [71]) and background genes using a one-sided Mann-Whitney U (MWU) test, finding that known PD genes are significantly altered more at all time points (*p-value ≤* 2.65*e^−^*^02^) and confirming that “gene movement” can indeed be used for discovering new PD-related genes. Using the assumption from Mihajlovic *et al.* [61] that unknown PD-related genes are also characterized by higher “gene movement”, in the second step, MONFIT computes the “gene movements” for each time point, representing each gene with a vector of “gene movements” across all time points. In the third step, MONFIT computes the length of this vector as an indicator of the TGM, ranking the genes based on the magnitude of their TGMs, from largest to smallest TGM, and predicting novel PD-related genes by focusing on the genes with the highest TGMs. To obtain these genes, MONFIT computes the elbow point of the curve of ranked *TGM* values and thresholds the genes, keeping only genes with TGM larger than the elbow point resulting in 163 multi-omics gene predictions (Supplementary Figure 3A; Supplementary File 1). Importantly, MONFIT does not use annotations of PD genes for its prediction selection process. Instead, it utilizes these annotations solely to validate the applicability of its downstream mining method.

Therefore, to determine the PD relevance of our novel gene predictions, we validate them by performing an enrichment analysis in the sets of known PD genes obtained from DisGeNet (1,148 genes) [71], Gene4PD (2,195 genes) [51], and their union (2,989 genes), which are also expressed in our scRNA-seq data across all time points. We find a statistically significant enrichment for all sets of these known PD genes (Supplementary Figure 3B) and observe the highest enrichment for the union set of known PD genes (*p-value* = 3.55*e^−^*^04^), with 41.7% of our gene predictions belonging to this gene set. For a complementary validation of our predictions, we conduct an additional validation experiment by computing the co-occurrence of each predicted gene with the term “Parkinson’s disease” in PubMed publications and compare these with a background set of genes that are expressed across all time points, but not among the gene predictions. Applying a one-sided MWU test between the two co-occurrence distributions, we find that the co-occurrence distribution of our 163 gene predictions is significantly greater than the one of the background (*p-value* = 1.74*e^−^*^11^). These results demonstrate that the 163 gene predictions are significantly associated with PD and can be used to propose novel PD-related genes.

### 3.3 Pathway enrichment analysis of MONFIT gene predictions

To determine the biological roles of our identified 163 gene candidates and their relation to PD by molecular mechanisms, we perform a pathway enrichment analysis (Supplementary Section 2) in RPs [41] and GO-BPs [4] and investigate whether the enriched pathways are known in the literature to be implicated in PD.

Reactome pathway enrichment analysis reveals 59 significantly enriched pathways (Supplementary File 2). To summarize the enriched pathways and ease their interpretability, we classify the enriched pathways based on the hierarchy of RPs and take the top-level RP as the representative term. We observe the following top three representative Reactome pathways that summarize the most pathways (Supplementary File 2): *Metabolism of RNA*, *Disease* and *Metabolism of proteins*. In addition, we perform an enrichment analysis of the 163 gene predictions in GO-BP terms and find significant enrichment in 39 terms (Supplementary File 3), which are summarized with REVIGO [80] (Supplementary Section 3) and lead to the representative GO terms that include: *translation*, *rRNA processing* and *antimicrobial humoral immune response mediated by antimicrobial peptide*. From both analyses, we observe that the enriched pathways converge on disrupted protein machinery starting from aberrant metabolism of RNA, including rRNA processing and translation, which leads to the dysregulated metabolism of proteins, triggering an autoimmune response and cellular death, which we discuss in the following.

Altered *rRNA processing* in PD substantia nigra could be a response to preserve energy homeostasis by decreasing gene expression of ribosomal protein mRNAs. This, in turn, causes a decreased synthesis of ribosome subunits, which may lead to a negative feedback reaction that contributes to cell damage and neuronal loss [31]. It should be noted that 21.5% of our 163 PD gene predictions are ribosomal genes, indicating the important role of ribosomal synthesis in PD. *Translation*, a process downstream of *rRNA processing*, has been implicated in PD, as several PD-related proteins have been shown to intervene with this process. For example, *LRRK2* (a commonly mutated protein in PD) modulates the amount of eukaryotic initiation factors, disrupting the initial phases of mRNA translation and promoting PD development [19]. The same effect could be achieved through *PINK1* mutations, as they influence *LRRK2* levels [5]. However, while mutations in many PD genes affect translation in a way that promotes PD pathogenesis, others, such as *PINK1*, are shown to suppress translation, leading to negative regulation of translation as a neuroprotective mechanism to lower energy consumption under stress conditions [59]. Thus, the role of translation in PD should be investigated further. Aberrant translation could, in turn, dysregulate *Metabolism of proteins*. For example, PD is characterized by the misfolding of *α*-synuclein and the formation of the Lewy Bodies, resulting in increased neuronal toxicity and cell death [72]. In addition, disrupted protein folding of tubulin can induce toxic accumulation of misfolded tubulin heterodimers, modifying microtubule stability and leading to early cytoskeletal dysfunction that is suggested to underline PD pathogenesis [69]. Finally, *PINK1* mutations impair the ability of *PINK1* to promote parkin phosphorylation and activate parkin-mediated ubiquitin signaling, resulting in the negative regulation of this pathway [75].

Enrichment in *antimicrobial humoral immune response mediated by antimicrobial peptide* term is indicative of an immune system response. Numerous studies demonstrated that PD triggers an immune system response resulting in neuroinflammation that perpetuates the neurodegenerative process [81]. This is further supported with the enrichment of *Disease*-related pathways, which include SARS-CoV-1 and SARS-CoV-2. For example, the immune response triggered by the activation of the NF*κ*B signaling cascade leads to the production of proinflammatory cytokines characteristic for PD [26] and infectious diseases such as SARS-CoV-1 and SARS-CoV-2 [16, 35, 21].

These findings confirm the relevance of our predictions to PD and underscore their potential to guide future studies aimed at exploring the disease’s aetiology and devising novel therapeutic strategies that would ameliorate the effects of PD by targeting and regulating these molecular mechanisms. Specifically, the enrichments in *rRNA processing*, *Translation*, *Metabolism of proteins* and immune response-related pathways are indicators that early PD development may be driven by the dysregulated protein synthesis machinery that triggers a cascade of events culminating in misfolded and aggregated *α*-synuclein proteins which disrupt cellular homeostasis, triggering an autoimmune response causing neuronal death. This is in line with findings and the prevailing understanding that Parkinson’s disease is associated with autoimmune features [22]. Thus, these pathways may play a crucial role in the pathogenesis of PD, further highlighting the need to study the role of these mechanisms and their relationship in the context of PD.

### 3.4 Top gene predictions are associated with Parkinson’s disease

The results above show that our PD gene predictions are globally associated with PD. To evaluate our ranking approach and propose new PD-associated genes, we focus on the top 30 prioritized PD predictions and characterize their relationship with PD. We manually assess if they have been implicated in PD in the literature and find that 25 genes (83.33%) have a known association with PD (Table 1), further demonstrating that our methodology predicts and prioritizes PD-relevant genes. In addition, we discover that the five (*CENPF*, *CRABP1*, *TOP2A*, *TMSB10*, and *NASP*) gene predictions have not previously been linked to PD, but find evidence in the literature that indicates their potential involvement in PD and propose them as novel PD candidates.

**Table 1:** Validation of the top 30 MONFIT gene predictions. The table ranks genes according to their “total gene movement” (TGM), so that genes with the largest TGM are ranked at the top. Genes in bold have literature that supports their direct role in PD. The number in the Evidence field is the PubMed ID number of the study, showing why the prediction is relevant for PD. For two PubMed IDs separated by an arrow in the Evidence field, the study labeled with the first PubMed ID implicates the gene in a biological mechanism/function and the second one explains how the mechanism/function is associated with PD. Genes whose basic function described in GeneCards [78] is PD-related have been annotated with GeneCards in the Evidence field.

| Rank | Gene | Evidence |
| --- | --- | --- |
| 1 | **MALAT1** | 35870046 |
| 2 | **COL1A2** | 37108598 |
| 3 | **VIM** | 31699368 |
| 4 | **DLK1** | 36820885 |
| 5 | **LUC7L3** | 33148011 |
| 6 | **WSB1** | 27273569 |
| 7 | **STMN1** | 32082315 |
| 8 | CENPF | 34857915 → 24768991 |
| 9 | **HMGB2** | 25306442 |
| 10 | CRABP1 | 10506831 → 25798108 |
| 11 | **MAP2** | 11406314 |
| 12 | **GAP43** | 36820885 |
| 13 | TOP2A | GeneCards → 16600170 |
| 14 | **RTN4** | 34830564 |
| 15 | **MAP1B** | 31235589 |
| 16 | TMSB10 | GeneCards, 37348871 → 27600680 |
| 17 | **HSP90AA1** | 34919646, 34626793 |
| 18 | **HSP90AB1** | 35731466 |
| 19 | **ENO1** | 36768134 |
| 20 | **TTC3** | 37677864 |
| 21 | **PTMA** | 35126116 |
| 22 | **PEG10** | 35725899 |
| 23 | **GPC3** | 35027645 |
| 24 | **FTL** | 33507490 |
| 25 | NASP | GeneCards → 34626793 |
| 26 | **WLS** | 24108702 |
| 27 | **NREP** | 28965931 |
| 28 | **TUBA1B** | 33732455 |
| 29 | **AKAP9** | 23832570 |
| 30 | **TUBA1A** | 19052140 |

As the expression of *CENPF* regulates the phosphorylation of ERK [17] and phosphorylated ERK is observed within degenerating neurons of PD [85], *CENPF* could promote PD by affecting this process. *CRABP1* binds retinoic acid, regulating its bioavailability and metabolism [63]. Retinoic acid supports the development and functional maintenance of dopaminergic neurons in PD, and its deletion leads to the development of neurodegenerative diseases, including PD [28]. Therefore, altered *CRABP1* negatively affects the uptake of retinoic acid, directly implicating *CRABP1* in PD pathogenesis, which prompts additional investigations. *TOP2A* encodes an enzyme that controls and alters the topologic states of DNA during transcription by catalysing the transient breaking and rejoining of two strands of duplex DNA [78]. Hegde *et al.*[36] showed a significantly higher number of DNA strand breaks in the PD-midbrain than in the age-matched control. Hence, altered activity of *TOP2A* may disrupt healthy cellular functioning already at the genomic level, leading to the development of PD. *TMSB10* plays a vital role in cytoskeleton organization [78], whose response to Alzheimer’s disease (AD) may mitigate the central nervous system damage caused by this disease [84]. As PD and AD share many mechanisms of neurodegeneration [89], including cytoskeletal dysfunction [69], *TMSB10* represents an attractive new candidate for further studies on the relation of the cytoskeleton with both diseases. Finally, *NASP* forms a cytoplasmic complex with *HSP90AA1* (rank 16 on our list of predictions, Table 1) and stimulates *HSP90AA1* ATPase activity. *HSP90AA1* is a known PD-related protein whose inhibition was found to reduce the aggregation of *α*-synuclein and *α*-synuclein-induced toxicity in a cellular PD model [38]. Thus, *NASP* could be involved in the aetiology of PD, through its relationship with *HSP90AA1*.

In conclusion, the five proposed PD candidates participate in molecular mechanisms whose disruptions are intrinsically tied to PD and its progression, implying their role in the pathogenesis of this disease and warranting future studies.

### 3.5 Comparing PD gene predictions with DEGs and DAPs

To explore if MONFIT predicts PD-associated genes beyond the standard approaches based on differential analyses, we compare our predictions with differentially expressed genes (DEGs) and differentially abundant proteins (DAPs) obtained from Bernini et al. [8] (the original study of the data) independent single-omics analyses that we filter to include the most significant DEGs (502 genes) and DAPs (2,240 genes) (see Supplementary Section 6).

The comparison between our 163 gene predictions and the most significant DEGs and DAPs shows that 23 (14.1%) predictions are DEGs, 16 (9.8%) are DAPs, and 36 (22.1%) are both DEGs and DAPs (Figure 2A), indicating that beyond the 46% overlap with the original study, MONFIT prioritizes a different set of genes than the conventional approaches. In addition, from the five newly proposed PD genes (Section “Top gene predictions are associated with Parkinson’s disease”), one (*TOP2A*) is a DEG, two (*CENPF* and *CRABP1*) are both DEGs and DAPs, while the remaining two (*TMSB10*, *NASP*) are neither DEGs nor DAPs, further proving that MONFIT can predict new disease genes that would have remained undetected with standard analysis approaches. To better position our findings with respect to conventional DEG and DAP analyses, we perform enrichment analysis in RPs and GO-BPs of the set of 440 unique DEGs (i.e., DEGs that are not MONFIT predictions), the set of 2,188 unique DAPs (i.e., DAPs that are not MONFIT predictions) and the set of 85 unique MONFIT predictions (i.e., MONFIT predictions that are not DEGs or DAPs). We find that the unique DEGs, DAPs, and MONFIT predictions are significantly enriched in 12, 19, and 67 RPs, with DEGs and DAPs sharing five RPs (Figure 2B). By summarizing the enriched RPs using the Reactome hierarchy, we find that DEGs and DAPs are enriched in pathways, that belong to a *Transmission across chemical synapses* subgroup. The perturbation of these mechanisms caused by dysfunctional genes might represent early signs of disrupted ability to release dopamine from mDA neurons [20], leading to the appearance of motor symptoms characteristic of PD. DAPs are additionally associated with the *Metabolism of lipids*, specifically mitochondrial fatty acid beta-oxidation, and *Synthesis of DNA*, whose disruption has been observed in PD [13, 36]. From the 67 RPs enriched for the 85 unique MONFIT predictions, 53 include pathways enriched in the entire set of 163 MONFIT predictions, with pathways converging on *Metabolism of RNA*, *Disease* and *Metabolism of proteins*, which have already been discussed in the context of PD in Section “Pathway enrichment analysis of MONFIT gene predictions”. By focusing on GO-BPs, we find that the unique DEGs, DAPs, and MONFIT predictions are significantly enriched in 59, 0, and 33 GO-BPs (Figure 2C). To summarize the GO-BPs, we use REVIGO [80] (Supplementary Section 3) and observe that many GO-BPs enriched for unique DEGs describe processes involved in *axon guidance and genesis* and *chemical synaptic transmission*. The abnormal functioning of axon guidance pathways can disrupt the modulation and maintenance of synaptic transmission leading to PD [54]. GO-BPs enriched for unique MONFIT predictions are related with *rRNA processing* and *Metabolism of proteins*. The alterations in these two basic processes may be precursor events driving the changes in the axon-guidance pathways, the loss of synaptic connectivity, and, ultimately, PD. Overall, these results indicate that genes predicted only by MONFIT point to pathways that underline PD pathogenesis, complementing the knowledge that could be gained by only studying mechanisms highlighted by DEGs and DAPs, which tend to converge on pathways that are a consequence of the disease.

**Figure 2:**
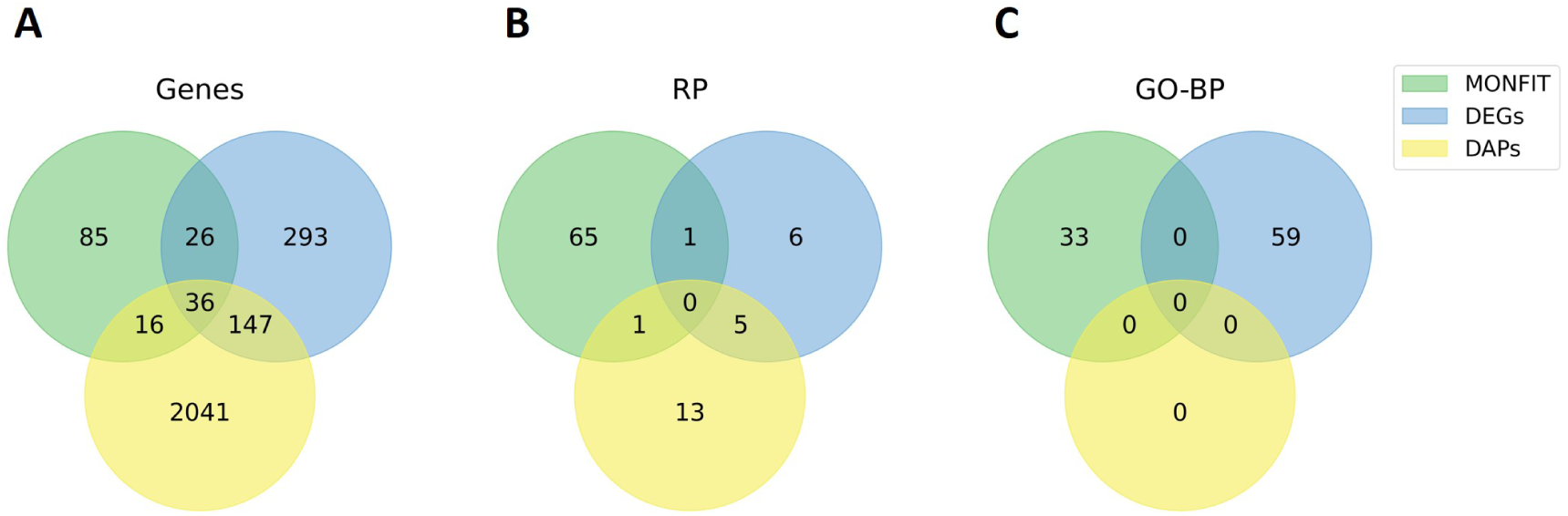
Comparing genes predictions obtained from MONFIT with DEGs and DAPs obtained from Bernini et al. [8]. **(A)** Overlap of MONFIT gene predictions with DEGs and DAPs by analyzing single-omics data. **(B)** For 85 genes uniquely predicted by MONFIT, 440 DEGs that are not predicted by MONFIT and 2,188 DAPs, we perform an enrichment analysis in RPs and determine the overlap between the enriched terms. **(C)** Same as **(B)**, but for GO-BPs.

Furthermore, all three analyses identify 36 common genes, implying that these multi-validated genes play important roles in the development of PD. Although 27 genes (Supplementary Table 4) have been implicated in PD, the studies mainly report the altered expression of these genes or altered abundance of their protein products in PD samples compared to control and only rarely describe the mechanisms by which they contribute to the aetiology or progression of PD. This provides the opportunity for future studies to investigate the exact roles of these genes in PD. From the remaining nine genes, the relationship between two genes (*CENPF* and *CRABP1*) and PD has been discussed in Section “Top gene predictions are associated with Parkinson’s disease”. In addition, while a direct association between PD and seven of the multi-validated genes (*IGFBP5*, *SMC4*, *ATAD2*, *SULF1*, *ANP32E*, *PRIM1*, and *CENPH*) has not yet been observed, the molecular mechanisms they participate in make them interesting new candidates in the pathophysiology of PD. *IGFBP5* is a binding protein that enables insulin-like growth factor I (IGF-1) binding activity [78], which plays a crucial role in neurodevelopment and apoptosis [44]. Multiple evidence points to a clear association of IGF-1 with the development of PD [15]. For example, IGF-1 levels are elevated in PD patients at the onset of the disease [15], suggesting that *IGFBP5* could induce or promote PD by regulating the activity of IGF-1. *SMC4* is a positive regulator of the inflammatory innate immune response that leads to the production of proinflammatory cytokines [87], which is widely observed in PD patients and is critical for the propagation of brain inflammation and the PD neurodegeneration [57]. Determining the detailed involvement of *SMC4* in PD-related immune response could provide a better understanding of the disease process, possibly providing opportunities for treatment. *ATAD2* mediates DNA replication, DNA repair, and transcription [40], while *PRIM1* is involved in DNA replication and its initiation [78]. DNA damage and the impairment of DNA repair systems are suggested to lead to increased alpha-synuclein phosphorylation, which facilitates the aggregation and toxicity of Lewy bodies typical for PD [82] and puts additional stress on the cell, causing further DNA damage [62] and disrupted DNA replication and transcription, thereby creating an escalating feedback loop mechanism. Therefore, *ATAD2* and *PRIM1* may play important roles in PD pathogenesis that deserve further studies. *SULF1* regulates vascular endothelial growth factor signaling, which has shown neuroprotective effects on dopaminergic neurons in a rat model of PD, implying that any perturbations to *SULF1* may facilitate PD development by disrupting vascular endothelial growth factor signaling. *ANP32E* inhibits the activity of protein *PP2A* [78]. The activity of *PP2A* decreases with the loss of *PINK1* function in PD[91], suggesting that the combined effect of *PINK1* and *ANP32E* promote PD, prompting the need for studying the relationship between *ANP32E* and PD. Finally, *CENPH* participates in signaling by Rho GTPases pathway [78], which has been connected with neurodegeneration in PD [3]. This indicates that dysregulated *CENPH* could contribute to PD pathogenesis by disrupting Rho GTPases signaling.

The results demonstrate that joint and integrative analysis of different omics data provides a new and valuable perspective on the development of PD that could not have been uncovered through independent analysis of different data types. In addition, we show that MONFIT complements standard differential analysis approaches. The convergence of multiple analyses on a common set of genes implies their significance in PD, calling for more extensive studies to unravel their involvement in PD and potential intervention opportunities.

### 3.6 MONFIT gene predictions and their relationship with PINK1 in PPI network

To assess if MONFIT produces PD gene predictions that are relevant to the specific PD subtype featuring a *PINK1* mutation, we analyze the connectivity between our predictions and the *PINK1* gene in the PPI network by measuring the shortest path lengths of our 163 predictions to *PINK1* and the density of the PPI subgraph that the predictions form with *PINK1*. The intuition behind this approach is based on the guilt-by-association principle, where disease genes are considered to be close to each other in a molecular network and are likely to participate in the same functional modules [67]. In addition, we determine if our multi-omics predictions are more associated with the *PINK1* subtype of PD by evaluating if they are closer to the *PINK1* gene in the PPI network or form a more connected PPI subgraph than DEGs, or DAPs, obtained from Bernini et al.[8] and described in the previous Section “Comparing PD gene predictions with DEGs and DAPs”. To analyze the PPI network relevant to our PD data, we generate the subgraph of the PPI network obtained from BioGrid induced by the genes expressed across all cell conditions (15,821). In this data-specific PPI network, we measure the shortest path lengths of MONFIT gene predictions, DEGs, DAPs and the background (genes in the PPI subgraph that are not in the former three gene sets) to the *PINK1* gene obtaining four shortest paths distributions. From the identified DEGs and DAPs (Section “Comparing PD gene predictions with DEGs and DAPs”), we consider only those that are present in the data-specific PPI subgraph, resulting in 394 DEGs and 2,119 DAPs. A simple measure of the average shortest path length (1.85) indicates that MONFIT gene predictions are closer to the *PINK1* gene in the PPI network than DEGs (2.18), DAPs (2.1) or background (2.24). We further compare the shortest path length distributions using a one-sided MWU test. When comparing distributions of the MONFIT gene predictions, DEGs, or DAPs with the background, we observe that all three gene sets are statically significantly closer to the *PINK1* gene than the background (*p-value ≤* 9.67*e^−^*^03^, Figure 3), indicating that all gene sets are indeed specific to the *PINK1* subtype of PD. By comparing our gene predictions’ shortest path length distribution with those of DEGs or DAPs, we also observe that our multi-omics gene predictions are closer to the *PINK1* gene than genes prioritized by single-omics analyses (*p-values ≤* 7.25*e^−^*^11^). In addition, 38 (23.3%) MONFIT gene predictions are direct neighbors of *PINK1* in the PPI network, which is statistically significant, as confirmed by performing a hypergeometric test (*p-value ≤* 2.38*e^−^*^20^). This is in contrast to only 4.9% of DEGs and 4.5% of DAPs that are *PINK1*’s direct neighbours.

**Figure 3:**
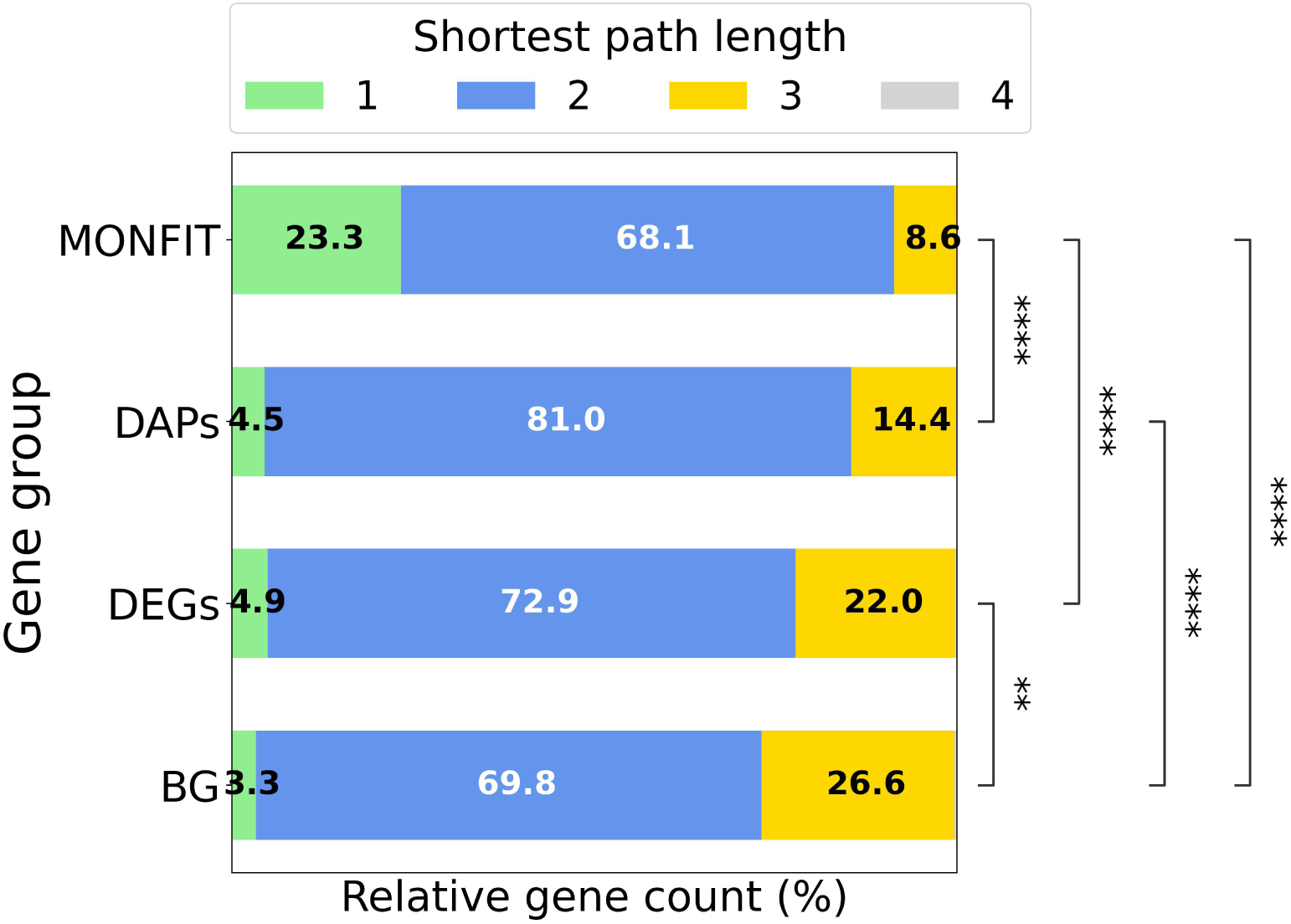
Shortest path lengths to *PINK1* in the PPI networks of MONFIT gene prediction, DEGs, DAPs and background (BG) genes (genes expressed in at least one cell condition, not including the gene predictions, DEGs and DAPs). Different patches of one gene set are proportions of genes that are 1, 2, 3 or 4 hops away from *PINK1*. The percentages of genes that are 4 hops away from *PINK1* is *≤* 1%, and their labels are not shown in the plot. We compare the shortest path length distributions using a one-sided Mann Whitney-U test. The *p-values* are adjusted for multiple hypothesis testing using the Benjamini-Hochberg procedure [7]. The *p-values* are indicated with stars for each comparison, so that: **\*\*\*\*** (*p ≤* 0.0001), **\*\*\*** (0.0001 *< p ≤* 0.001), **\*\*** (0.001 *< p ≤* 0.01), **\*** (0.01 *< p ≤* 0.05), and **ns** (*p >* 0.05, not significant).

These results suggest that DEGs and DAPs are more indicative of downstream biological changes that are a consequence of an initial cellular response to the *PINK1* mutation, when compared to MONFIT that uncovers genes that are closer to *PINK1* in the PPI network, thereby highlighting potential causalities and therefore better intervention points.

To evaluate the global connectivity of the PPI subgraph that *PINK1* forms with 163 MONFIT PD gene predictions, we perform sampling with replacement experiment across 10,000 runs and measure the density of the subgraph induced by our gene predictions in relation to *PINK1* and compare the density to a control PPI subgraph spanned by *PINK1* and 163 randomly chosen genes from the data-specific PPI. We find that the density of the PPI subgraph induced by our predictions and *PINK1* (5.04*e^−^*^02^) is statistically significantly higher than random (*p-value* = 9.99*e^−^*^05^), indicating that our predictions are globally well connected with *PINK1*. This might suggest that MONFIT PD gene predictions participate in molecular mechanisms that are directly disrupted by the *PINK1* mutation, triggering a cascade of events that lead to the development of PD. To further investigate if our predictions are more specific to the *PINK1* subtype of PD than the top prioritized DEGs or DAPs, we observe how well the PPI subgraphs induced by these gene sets are connected together with *PINK1*. We find that the PPI subgraph of DAPs, but not DEGs, with *PINK1* are statically significantly well connected by performing sampling with replacement (*p-value* = 9.99*e^−^*^05^) and observe that the densities (5.05*e^−^*^03^ for DEGs and 6.39*e^−^*^03^ for DAPs PPI subgraphs with *PINK1*) are smaller by an order of a magnitude than the density of the PPI subgraph formed by our MONFIT gene predictions and *PINK1*. This shows that our gene predictions are more connected with *PINK1* in the PPI network, indicating that our multi-omics integration methodology predicts genes that are more closely related to the PD subtype caused by *PINK1* mutation, compared with the genes prioritized according to the analysis of single omics data.

Overall, these results demonstrate the power of MONFIT to predict genes relevant to the specific PD subtype characterized by a *PINK1* mutation. Moreover, we demonstrate that our multi-omics gene predictions are more closely related to the *PINK1* gene, forming a denser PPI subgraph than the genes prioritized by standard single-omics differential analyses, suggesting that MONFIT predicts genes that are focused on disease-triggering events which are more indicative of disease causality.

### 3.7 Drug-repurposing candidates for PD

Existing therapeutic options for PD aim to reduce the severity of the symptoms, and there are no treatment options that stop the progression of the disease. Having uncovered a set of PD-associated genes and their related molecular pathways that potentially indicate where things first go awry in a cell line carrying a *PINK1* mutation causing PD, we rely on a drug-repurposing strategy to identify new treatment strategies for PD. To propose drug-repurposing options, we check for drug-target interactions of our 163 MONFIT gene predictions against a set of 2,519 approved and investigational drugs from DrugBank v5.1.10 [88] and find that 37 (22.3%) predictions are known drug targets of 94 drugs. We perform enrichment analysis (Supplementary Section 2) of our 163 PD gene predictions in the collected set of drug-target interactions by treating drugs as annotations and the interactions as indicators by which drugs target the corresponding genes. The analysis identifies 57 drugs that significantly target our gene candidates compared to background genes (genes expressed across all cell conditions), which we refer to as enriched drugs. From the *p-value*-ranked drugs, we discuss in the following the potential therapeutic effects on PD for the top 7 most significantly enriched drugs that target MONFIT gene candidates participating in pathways of the protein synthesis machinery and immune response (Section “Pathway enrichment analysis of MONFIT gene predictions” and Figure 4).

**Figure 4:**
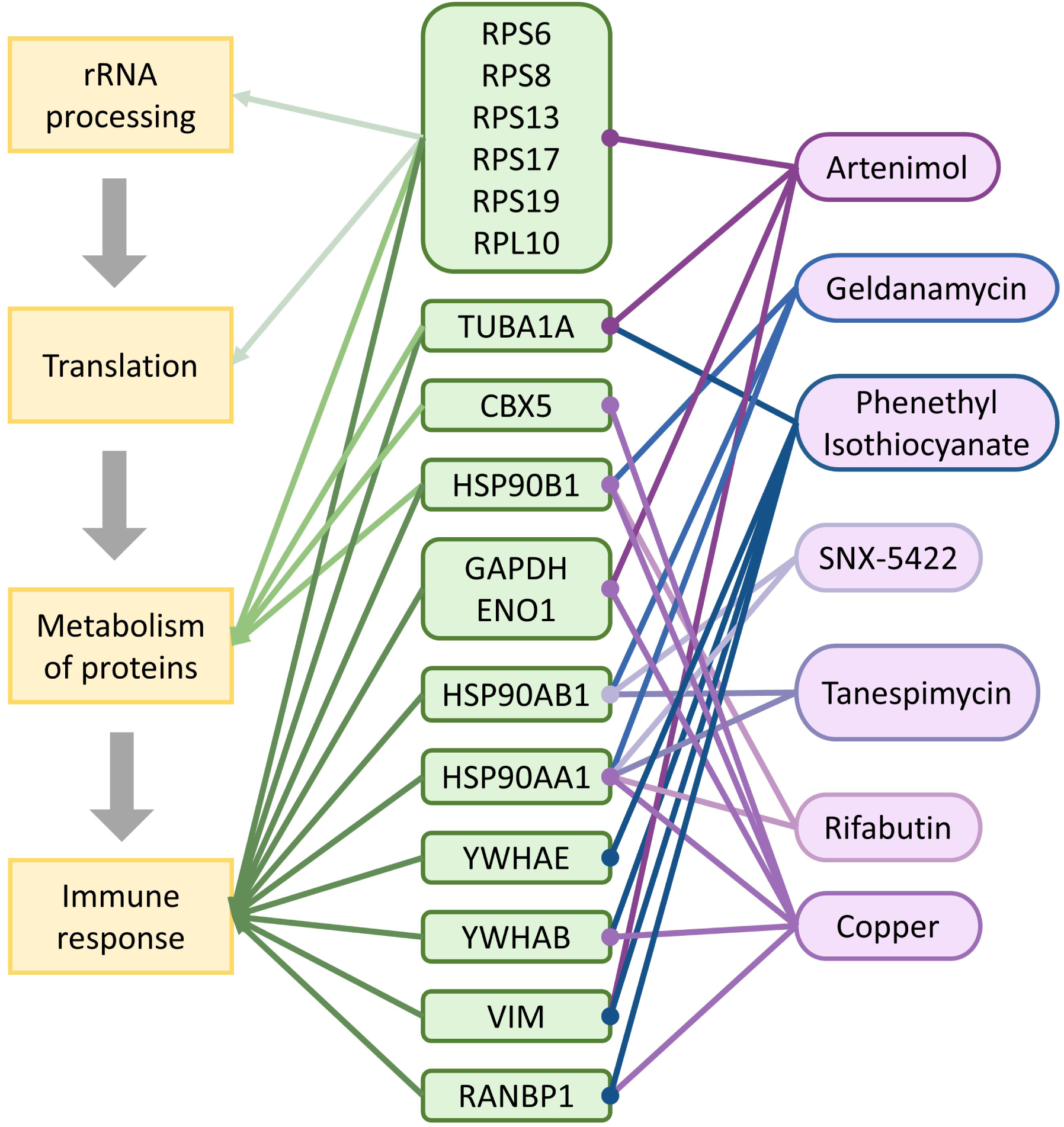
Diagram of drug-repurposing options for PD, targeting MONFIT gene predictions that participate in mechanisms central for PD progression that converge on disrupted protein synthesis machinery, followed by an immune response. Purple labels denote drugs, green labels represent genes, and yellow labels are molecular pathways. Dark grey arrows indicate the influence of one pathway on another. Green arrows with the same shades indicate which genes participate in one mechanism. Purple/blue lines with the same shades indicate which genes are targets of one specific drug.

Artenimol (i.e., dihydroartemisinin), the top enriched candidate (*p-value*= 3.86*e^−^*^10^), is targeting 14 gene predictions (six of which code for ribosomal proteins) that play different roles across all pathways of the protein synthesis machinery and immune response (Figure 4). Artenimol is an antimalarial agent [88], which could be used to treat PD because it has been shown to modulate the PI3K/AKT signalling pathway, thereby reducing neuroinflammation in mice [30]. PI3K/AKT signalling pathway is implicated in PD pathogenesis [55] and is a well-known regulator of ribosome biogenesis, which is decreased in PD [31]. Therefore, Artenimol could exert neuroprotective effects and rescue PD phenotype by regulating the disrupted PI3K/AKT signalling pathway, reestablishing normal rates of ribosome biosynthesis and reducing neuroinflammation. This evidence suggests that Artenimol might be a very attractive drug-repurposing candidate for PD the potential of which to modulate protein synthesis, metabolism, and immune response should be investigated in future studies.

Our study revealed Geldanamycin, a second most significantly enriched drug (*p-value*= 1.25*e^−^*^04^), whose mechanism of action is exerted by inhibiting the function of *HSP90* genes. Geldanamycin, which targets three gene predictions and belongs to the ansamycin group of antibiotics, can reduce *α*-synucelin aggregation by binding to and inhibiting the function of the *HSP90B1*, which promotes proteasomal degradation, thereby protecting against dopaminergic loss in PD. However, its hepatotoxicity, poor solubility and poor brain permeability prevent its use in a clinical setting [76]. Other studies suggest that Geldanamycin derivatives could overcome these problems, offering promising compounds worthy of additional studies [76, 90].

The third most significantly enriched drug (*p-value*= 1.49*e^−^*^02^) is an antioxidant drug phenethyl isothiocyanate, targeting six predictions, that was tested in an in vitro model for its potential to mitigate the oxidative stress involved in the pathogenesis of PD and reduce neuroinflammation [48]. Although in vitro experiments reported a moderate ability to counteract the production of reactive oxygen species causing oxidative stress [48], more studies are necessary to determine the neuroprotective capacities of this drug.

Similarly to Geldanamycin, Tanespimycin and Rifabutin (rank 4; *p-value*= 2.6*e^−^*^03^) are also a member of ansamycin antibiotics that inhibits *HSP90* thereby attenuating *α*-toxicity. However, Tanespimycin’s use in neurodegenerative diseases remains limited because of poor blood–brain-barrier permeability [27]. On the other hand, Rifabutin, reduces *α*-synuclein oligomers ameliorating dopaminergic neurodegeneration in the rat model of PD. Moreover, because Rifabutin can penetrate the blood-brain barrier, it is a very attractive drug-repurposing option for PD, presenting a viable drug choice for consideration in human clinical trials [18]. The last *HSP90* inhibitor on our list of most enriched drugs is SNX-5422 (rank 4; *p-value*= 2.6*e^−^*^03^), a water-soluble and orally bioavailable prodrug of SNX-2112 that has broad applicability across a wide range of cancers by inhibiting tumour growth [88]. It has also been investigated for treating SARS-COV-2 infection, effectively suppressing it by mitigating the virus-induced inflammation [33]. The anti-inflammatory characteristics of SNX-5422 and its capability to penetrate the blood-brain barrier make SNX-5422 a promising therapeutic strategy for treating PD. Tanespimycin, Rifabutin, and SNX-5422 are the fourth-ranked drugs as they all target two genes each and have the same enrichment *p-values*.

Finally, copper is the fifth most significantly enriched drug (*p-value*= 2.47*e^−^*^05^), targeting nine of our gene predictions that participate in pathways of protein metabolism, and immune response. Interestingly, both excessive and deficient levels of copper contribute to PD by increasing oxidative stress, leading to cell damage [10]. To address low copper levels in PD patients, a phase 1 clinical trial (NCT03204929) using a copper-delivering molecule *Cu*^+2^-ATSM, has been completed, but the results have not yet been published. More studies are necessary to investigate copper as a potential PD-modifying drug rescuing copper homeostasis, thereby regulating a chain of mechanisms whose dysregulation is implicated in PD (Figure 4).

In summary, our analysis provides promising drug candidates for disease-modifying therapeutics for PD. Future studies are needed to investigate their potential in treating PD to uncover agents that could be further tested in human clinical trials. Although some of our most enriched drugs are not clinically applicable, they could be replaced with more clinically relevant candidates with similar mechanisms of action. Our findings point to a similar set of drug candidates as the recent study aimed at identifying new therapeutics for PD [83], further emphasizing the potential of these drugs in treating PD and demonstrating the power of MONFIT to prioritize genes beyond classical DEG and DAP analyses for the identification of new intervention points in PD.

## 4 CONCLUSION

The complexity of PD requires new integration methods capable of exploiting time-resolved multi-omics data at bulk and single-cell levels, together with prior knowledge in molecular interaction networks, to investigate how PD-causing mutations alter gene expression, resulting in modified proteomics profiles and dysregulated metabolism.

In this work, we present MONFIT to integrate multi-omics data with prior knowledge in the molecular interaction networks of PPI, MI, GI, and COEX. We apply MONFIT to time-series measurements of scRNA-seq data, bulk proteomics, and bulk metabolomics of a PD patient-derived iPSC cell line harboring a *PINK1* mutation and a control line. For this purpose, we construct data-driven PPI and MI networks by incorporating the bulk proteomics with PPI from BioGRID and bulk metabolomics with MI from KEGG, by weighting the network edges with the measured omics data. To ensure gene embeddings that effectively capture the molecular mechanisms related to PD, we conduct a grid-search analysis to determine the optimal combination of input data. Next, we use the notion of “gene movement” to compare the PD and control gene embeddings across time points and identify genes with the highest TGM across all time points leading to 163 gene predictions that are significantly associated with PD in literature. By performing enrichment analysis in RPs and GO-BPs, we demonstrate that our predictions participate in pathways with documented association with PD, highlighting multiple pathways as potential intervention points for halting PD. In addition, we manually validate the top 30 predictions in the literature to discover that 25 genes are associated with PD and provide compelling evidence elucidating the role of the remaining five genes.

In the subsequent comparison of the MONFIT gene candidates with DEGs and DAPs, we demonstrate that our integrative multi-omics methodological pipeline goes beyond standard differential analysis approaches of single-omics data by uncovering PD-relevant genes that would otherwise remain undetected and are indicative of PD pathogenesis. This is further supported by studying the connectivity of MONFIT gene predictions, DEGs and DAPs with *PINK1* in the PPI network, where we see that our predictions are more closely connected with *PINK1* than DEGs or DAPs. This implies that MONFIT reveals genes that are pertinent not just for PD in general but for a specific subtype of PD caused by a *PINK1* mutation and that they are potential causalities and, therefore, better intervention points. Finally, we check if our predictions are drug targets to propose future therapeutic strategies based on drug repurposing.

To our knowledge, MONFIT is so far the only multi-omics integrative approach designed to fuse time-series scRNA-seq, bulk proteomics, and metabolomics data with molecular interaction networks to uncover novel disease-associated genes beyond traditional differential expression (or abundance) analyses. This approach emphasizes mechanisms related to disease pathogenesis and proposes treatment strategies based on drug repurposing. While we designed and applied MONFIT in the context of PD, it is a generic method and could be modified to accommodate data from tissue samples and other data types, such as epigenetics. In addition, MONFIT could be extended to incorporate drug-target interactions in the integration process by embedding drugs and genes in the same space. This opens avenues for identifying novel treatment strategies by discovering previously unidentified drug-target interactions, surpassing the limitations of drug repurposing solely based on known drug-target associations.

## Data Availability

All original code and reproducibility materials are available at 10.5281/zenodo.11396807 [60]. Note that this paper analyzes existing, publicly available data. Any additional information required to reanalyze the data reported in this paper is available from the lead contact upon request.

