## Supplementary Information for "MONFIT: Multi-omics factorization-based integration of time-series data sheds light on Parkinson’s disease"

June 3, 2024

#### 1 Robustness of NetSC-NMTF gene embeddings to the number of dimensions and weighting factors

To assess the robustness of our results, we perform sensitivity analyses to determine how i) the number of dimensions and ii) different combinations of weighting factors of the input matrices used for producing the lower-dimensional matrices with NetSC-NMTF influence the organization of gene embeddings, $U$. We measure the robustness of the integration framework to these parameters by using $U$ gene embeddings to compute $GMMs$ and compare them across different parameter configurations. We use $GMMs$ for comparison because they are the basis from which we obtain our MONFIT gene predictions.

To investigate how the number of dimensions used for producing the lower-dimensional matrices with NetSC-NMTF influences the organization of gene embeddings, we perform data integration varying the number of dimensions used for producing embeddings, $k_1$ and $k_2$. For each cell condition, we apply a grid search approach centered on $k_{1_o}$ and $k_{2_o}$ values, used to obtain gene embeddings presented in the manuscript, so that $k_1 \in \{k_{1_o} - 30, k_{1_o} - 15, k_{1_o}, k_{1_o} + 15, k_{1_o} + 30\}$ and $k_2 \in \{k_{2_o} - 10, k_{2_o} - 5, k_{2_o}, k_{2_o} + 5, k_{2_o} + 10\}$. These parameter ranges aim to explore the immediate vicinity of the initial parameter choices, ensuring a focused investigation on the impact of the organization of gene embeddings. For each cell condition, we compute a $GMM$ for each combination of $k_1$ and $k_2$ and evaluate the agreement between the $GMMs$ in an all-to-all manner by computing the root mean square error (RMSE) between all pairs of $GMMs$. Then, we average the RMSEs across all $GMM$ comparisons and across all cell conditions and observe a small average RMSE $8.38e^{-06} \pm 3.08e^{-06}$, demonstrating that our NetSC-NMTF integration framework is robust to the choice of dimensionality of embedding space.

We determine how the weighting factors ($w_{PPI}$, $w_{COEX}$, $w_{GI}$, $w_{MI}$, $w_E$), which scale the corresponding input matrices (PPI, COEX, GI, MI, and E) before applying NetSC-NMTF, affect the organization of the obtained gene embeddings. Thus, we compare the gene embeddings of each cell condition by analyzing the top 100 ranked combinations out of the 768 generated by varying $w_{PPI,COEX,GI,MI}$ across $\{0, 0.1, 1, 10\}^4$ and $w_E$ across $\{0.1, 1, 10\}$. The highest ranked weighting factor combinations are those that lead to gene embeddings that best capture the molecular mechanisms associated with PD (Section "Finding the optimal combination of data for integration"). For each cell condition, we evaluate the agreement between gene embeddings associated with different configurations of weighting factors by comparing their $GMMs$ in an all-to-all manner by computing the RMSE between all pairs of $GMMs$. Finally, we compute the average RMSE for all comparisons made with $GMM$ across all cell conditions, resulting in a small average RMSE of $1.99e^{-05} \pm 9.32e^{-06}$. This shows that our NetSC-NMTF integration framework is robust to the choice of the weighting factor combination.

Overall, these experiments demonstrate the robustness of our MONFIT integration model.

#### 2 Enrichment analysis

To perform enrichment analysis in a set of annotations, we apply the same method used in Mihajlovic et al. [6], where we compute the probability, $p$ (i.e., *p-value*), that an annotation is enriched in a set (or a cluster) of genes by using a hypergeometric test (i.e., sampling without replacement strategy) [8]:

$$p = 1 - \sum_{i=0}^{X-1} \binom{K}{i} \binom{M-K}{N-i} / \binom{M}{N} \quad (1)$$

where $N$ is the number of annotated genes in the set (cluster), $X$ is the number of genes in the set (cluster) annotated with the given annotation, $M$ is the number of annotated genes in the corresponding background set of genes, and $K$ is the number of genes in the corresponding background set of genes annotated with the given annotation. We adjust the *p-values* to account for multiple hypothesis testing using the Benjamini-Hochberg procedure [1]. All annotations with an adjusted *p-value* $\leq 0.05$ are considered to be statistically significantly enriched.

#### 3 Summarizing the results of enrichment analysis in Gene Ontology Biological Processes

To determine the biological roles of MONFIT gene predictions, we perform a pathway enrichment analysis in Gene Ontology Biological Processes (GO-BP) terms, which may result in long lists of enriched annotations that are difficult to interpret.

To summarize the enrichment analysis in GO-BP terms, we apply REVIGO. REVIGO summarizes a list of GO terms by clustering them based on a semantic similarity measure thresholded at a user-specified cutoff value. The process can be guided with the user-provided enrichment *p-values* to select the most representative term. To generate the most semantically diverse list of representative GO terms, without removing the terms that have no strong statistical support for their redundancy, we set the cutoff value at 0.5, corresponding to the "small" list size.

### 4 Bulk metabolomics and proteomics data are informative for predicting novel PD-associated genes

To determine if integrating bulk omics data contributes to predicting novel PD genes, we apply MONFIT on PD and control cell conditions at D8, D18, D25, D32 and D37 (time points which we use to obtain 163 PD gene predictions presented in the main manuscript) without including the bulk data during integration. For this purpose, we neither weigh the edges in the PPI networks with proteomics data nor the edges in the MI networks with metabolomics data, leading to binary adjacency matrices for PPI and MI networks, where one indicates interaction and zero indicates that there is no interaction between the nodes. To ensure that the difference in the obtained gene embeddings arises solely from the absence of bulk proteomics and metabolomics data during integration, we apply the same weighting factors to the input matrices that we use for integrating all multi-omics data (presented in the main), so that $w_{PPI} = 1$, $w_{COEX} = 1$, $w_{GI} = 0$, $w_{MI} = 10$ and $w_E = 10$.

To evaluate if the resulting condition-specific gene embeddings are functionally organized to capture PD-relevant mechanisms, we cluster the gene embeddings of each cell condition with k-means clustering (the number of clusters is the same as the dimension of the embedding space) and perform enrichment analysis (hypergeometric test; Supplementary Section 2) in PD-related pathways from PD map [3], measuring the percentage of clusters and genes with enriched annotations. We average the percentage of clusters and genes with enriched annotations across all cell conditions and find 18.2% of genes and 23.9% of clusters with enriched PD map pathways, indicating that the embeddings obtained by not including bulk data during integration also capture biological features of PD. Next, we analyze the "gene movements" obtained from step 2 of MONFIT to investigate when gene embeddings of PD genes (obtained from DisGeNet [7]) display differences between PD and control cell conditions. We compute the "gene movements" of PD and background genes, comparing them with a one-sided Mann-Whitney U (MWU) test and observing that the movement of PD genes is significantly higher between PD and control cell conditions at three time points (day (D)8, D18 and D37; $p\text{-}value \leq 1.34e^{-03}$), in contrast to all five time points when bulk data is included in the integration. This indicates that bulk data contains valuable biological information on PD that complements the signal captured by SC data and knowledge-based molecular networks.

Finally, we apply the third step of MONFIT using the "gene movements" of time points D8, D18 and D37 of cellular development (i.e., time points where "gene movement" of known PD genes is significantly higher than background) to compute "total gene movement" (TGM) and rank the genes according to the highest TGM. By focusing on the genes with the highest TGMs (Section "MONFIT pipeline"), MONFIT predicts 119 genes. We validate the PD relevance of MONFIT gene predictions obtained by not including bulk data during integration in the literature by comparing their co-occurrence with the term "Parkinson's disease" in PubMed publications against the background, applying a one-sided MWU test ($p\text{-}value = 3.16e^{-13}$). Additionally, we perform an enrichment analysis in the sets of known PD genes obtained from DisGeNet (1,170 genes) [7], Gene4PD (2,244 genes) [5], and their union (3,052 genes) (all genes are expressed in our scRNA-seq data across time points D8, D18 and D37), finding a statistically significant enrichment for all sets of these known PD genes ($p$-$value \leq 1.38e^{-04}$). Therefore, we conclude that MONFIT predicts PD-relevant genes without the information contained in the bulk proteomics and metabolomics data.

However, we observe that excluding bulk data during integration results in a 27% decrease in predicted genes – from 163 (presented in the main that are based on integrating bulk data as well) to 119 gene predictions, with an overlap of 103 genes. In addition, by integrating bulk omics data, MONFIT predicts 60 genes that would otherwise remain undetected. These genes participate in pathways related to translation, metabolism of RNA, and infectious disease, whose importance for PD has been discussed in Section "Pathway enrichment analysis of MONFIT gene predictions". This suggests that integrating bulk omics data contributes to predicting PD-associated genes whose multi-omics profiles support their involvement in PD, implicating that such genes are relevant for the disease progression and could potentially lead to the pathogenesis of PD.

These results show that integrating bulk omics with SC data and molecular networks contributes to predicting novel PD genes.

### 5 Comparing the downstream method of MONFIT with the downstream method presented in Mihajlovic *et al.* [6]

To demonstrate that the downstream mining method employed by MONFIT improves the prediction of novel PD genes, we compare them with predictions obtained by the downstream analysis approach that inspired MONFIT's pipeline, which is presented in Mihajlovic *et al.* [6].

We analyze the gene embeddings produced by MONFIT (see Section "Finding the optimal combination of data for integration"), following the 2-step downstream analysis method from Mihajlovic *et al.* [6]. First, we cluster the gene embeddings encoded in $G_1$ matrices of each PD cell condition and perform an enrichment analysis in known PD genes from DisGeNet [7]. From the significantly enriched clusters, we retain those genes that are not PD genes in DisGeNet, obtaining ***Stage-specific PD predictions***. Second, we intersect all ***Stage-specific PD predictions*** to define ***Core PD predictions***, which we prioritize by computing the average "movement" of each prediction across all time points and ranking them according to the largest average "movement". Following this approach results in 411 ***Core PD predictions*** which overlap in only one gene with the 163 MONFIT PD predictions presented in the main.

We quantify the association of 411 ***Core PD predictions*** with PD in the literature by using an automated PubMed publication search to count the co-occurrence of each ***Core PD prediction*** and the background genes (genes expressed across all cell conditions that are not ***Core PD predictions***) with the term "Parkinson's disease" in PubMed publications. To measure if ***Core PD predictions*** are significantly more co-occurring with PD in the literature, we perform a one-sided MWU test (with a significance level of 0.05) between the co-occurrence distributions of the set of predictions and the background. We observe that the 411 ***Core PD predictions*** are co-occurring more than the background in PD-related publications ($p$-$value = 3.99e^{-03}$), as the predictions obtained with the downstream analysis of MONFIT. However, by performing an enrichment analysis of ***Core PD predictions*** in known PD genes from Gene4PD [5], we show that they are not enriched in this set of known PD genes (non-significant $p$-$value > 0.05$).

To investigate if ***Core PD predictions*** are associated with the specific PD subtype harboring a *PINK1* mutation, we analyze the PPI subgraph that these predictions form with *PINK1*. First, we measure the shortest path lengths of the 411 ***Core PD predictions*** to *PINK1* and compare them with those of the 163 MONFIT gene predictions. We observe that the average shortest path length of ***Core PD predictions*** (2.06) is larger than the one of MONFIT gene predictions (1.85). By comparing the two distributions of the shortest path lengths using a one-sided MWU test, we observe that the distribution of the shortest path lengths of MONFIT gene predictions is statistically significantly smaller than the one of ***Core PD predictions*** ($p$-$value = 4.35e^{-07}$; see Supplementary Figure 4). Additionally, only 5.4% of ***Core PD predictions*** are the first neighbors of *PINK1* in the PPI network, as opposed to 23.3% of MONFIT gene predictions. These results indicate that MONFIT predicts genes that are more relevant to the PD subtype caused by a *PINK1* mutation than the 2-step downstream method introduced in [6]. This is further supported by measuring the density of the PPI subgraph induced by ***Core PD predictions*** and *PINK1* ($9.99e^{-03}$) and observing that it is 5 times lower than the density of the PPI subgraph induced by MONFIT gene predictions and *PINK1* ($5.04e^{-02}$).

Overall, these results show that the downstream analysis of MONFIT produces predictions that are more relevant for PD than those obtained by employing the 2-step downstream analysis method from Mihajlovic *et al.* [6].

### 6 Obtaining the most significant differentially expressed genes and differentially abundant proteins

To explore if MONFIT predicts PD-associated genes beyond the standard approaches based on differential analyses, we compare our MONFIT gene predictions with differentially expressed genes (DEGs) and differentially abundant proteins (DAPs) from the original study of the data [2]. In the DAP table, the name of a protein corresponds to the name of the gene that codes for that protein. From Bernini *et al.* [2], we obtain all DEGs and DAPs whose fold change (FC) is greater than 0 in at least one-time point, taking into account days D8, D18, D25, D32 and D37, which we investigate in this study. Following the approach from Bernini *et al.* [2], we obtain the most significantly differentially expressed genes by filtering DEGs to include genes whose $FC > 0.5$, in at least one-time point, resulting in 502 genes. We perform the same filtering procedure on DAPs, resulting in 2,240 genes.

### 7 Supplementary Tables

| CC | PPI | | COEX | | MI | | GI | |
|---|---|---|---|---|---|---|---|---|
| | #nodes | #edges | #nodes | #edges | #nodes | #edges | #nodes | #edges |
| $C_{D8}$ | 13,851 | 379,078 | 12,087 | 1,191,230 | 2,156 | 105,632 | 4,922 | 13,365 |
| $C_{D18}$ | 14,106 | 385,174 | 12,267 | 1,203,102 | 2,194 | 109,189 | 4,959 | 13,501 |
| $C_{D25}$ | 14,143 | 392,108 | 12,311 | 1,205,719 | 2,196 | 109,046 | 4,955 | 13,454 |
| $C_{D32}$ | 14,115 | 372,677 | 12,226 | 1,200,812 | 2,184 | 107,770 | 4,948 | 13,377 |
| $C_{D37}$ | 13,085 | 332,387 | 11,535 | 1,153,654 | 2,043 | 91,751 | 4,769 | 11,456 |
| $PD_{D8}$ | 14,213 | 402,455 | 12,379 | 1,210,526 | 2,197 | 112,088 | 4,971 | 13,522 |
| $PD_{D18}$ | 13,817 | 393,848 | 12,075 | 1,190,984 | 2,155 | 106,194 | 4,892 | 13,384 |
| $PD_{D25}$ | 14,078 | 394,989 | 12,259 | 1,201,979 | 2,183 | 105,165 | 4,943 | 13,302 |
| $PD_{D32}$ | 12,558 | 330,846 | 11,117 | 1,116,413 | 1,966 | 84,440 | 4,632 | 12,582 |
| $PD_{D37}$ | 11,723 | 266,726 | 10,447 | 1,041,304 | 1,837 | 74,188 | 4,418 | 11,588 |

Supplementary Table 1: **Number of genes (#genes) and interactions (#edges) of each data-driven molecular network per cell condition (CC).** A cell condition is defined by a cell line at a specific time point (e.g., D8). A cell line harbouring a PINK1 mutation is referred to as PD and the control cell line as C.

| CC | #genes | #SCs |
|---|---|---|
| $C_{D8}$ | 13,851 | 2,788 |
| $C_{D18}$ | 14,106 | 2,757 |
| $C_{D25}$ | 14,143 | 2,713 |
| $C_{D32}$ | 14,115 | 2,767 |
| $C_{D37}$ | 13,085 | 1,571 |
| $PD_{D8}$ | 14,213 | 2,657 |
| $PD_{D18}$ | 13,817 | 2,783 |
| $PD_{D25}$ | 14,078 | 2,789 |
| $PD_{D32}$ | 12,558 | 2,380 |
| $PD_{D37}$ | 11,723 | 888 |

Supplementary Table 2: **Number of genes (#genes) and single cells (#SCs) of expression matrices for each cell condition (CC).** A cell condition is defined by a cell line at a specific time point (e.g., $C_{D8}$, for Control cell line at day 8). A cell line harbouring a PINK1 mutation is referred to as PD and the control cell line as C.

| CC | $k_1$ | $k_2$ |
| --- | --- | --- |
| $\mathrm{C_{D8}}$ | 83 | 37 |
| $\mathrm{C_{D18}}$ | 83 | 37 |
| $\mathrm{C_{D25}}$ | 84 | 36 |
| $\mathrm{C_{D32}}$ | 84 | 37 |
| $\mathrm{C_{D37}}$ | 80 | 28 |
| $\mathrm{PD_{D8}}$ | 84 | 36 |
| $\mathrm{PD_{D18}}$ | 82 | 37 |
| $\mathrm{PD_{D25}}$ | 83 | 37 |
| $\mathrm{PD_{D32}}$ | 79 | 34 |
| $\mathrm{PD_{D37}}$ | 76 | 21 |

Supplementary Table 3: **Dimension parameters $k_1$ and $k_2$ used for decomposing the input data of each cell condition (CC).** PD: Parkinson's disease; C: Control.

| Rank | Gene | Evidence |
|---|---|---|
| 2 | **COL1A2** | 37108598 |
| 4 | **DLK1** | 36820885 |
| 8 | CENPF | 34857915 → 24768991 |
| 10 | CRABP1 | 10506831 → 25798108 |
| 12 | **GAP43** | 36820885 |
| 23 | **GPC3** | 35027645 |
| 26 | **WLS** | 24108702 |
| 34 | **MDK** | 37667404 |
| 40 | IGFBP5 | 31479860 → 35173238 |
| 46 | **NEFM** | 36777638 |
| 48 | **TFPI2** | 27191603 |
| 49 | **TPX2** | 37674708 |
| 57 | **STMN2** | 31748532 |
| 70 | **CNTNAP2** | 25475535 |
| 79 | **NCAM1** | 24853996 |
| 82 | SMC4 | 29803706 → 34239490 |
| 86 | **PCSK1** | 32858884 |
| 89 | ATAD2 | 29148850 → 24860424, 30050065 |
| 90 | **BNIP3** | 27528605 |
| 96 | SULF1 | 19373441 → 15066146 |
| 114 | **PCLO** | 38457145 |
| 120 | **SYT4** | 34305570 |
| 122 | **SLIT2** | 19162339 |
| 124 | **OTX2** | 35644505 |
| 126 | **TPBG** | 34876581 |
| 127 | **P4HA1** | [9] |
| 128 | ANP32E | GeneCards → 29416594 |
| 134 | PRIM1 | GeneCards → 25303312 |
| 146 | **MT2A** | 37107269 |
| 147 | **TCF7L2** | 36869069 |
| 148 | **LDHA** | 34973450 |
| 151 | **TXNIP** | 28755477 |
| 152 | **NEFL** | 36777638 |
| 156 | CENPH | GeneCards → 32878220 |
| 159 | **SPARC** | [4] |
| 160 | **CSRP2** | 35027645 |

Supplementary Table 4: **Literature validation of the genes that are in the overlap between MONFIT gene predictions, DEGs, and DAPs from the original study of the longitudinal omics data [2]**. Genes in bold have literature that supports their direct role in PD. The table ranks genes according to their TGM across all time points, so that genes with the largest TGM are ranked at the top. The ID number in the evidence field is the PMID number of a study that shows why a prediction is relevant for PD. For studies where PMID is not available, we provide a citation. Genes whose basic function described in GeneCards [10] is PD-related have been annotated with GeneCards in the Evidence field. For two PMIDs separated by an arrow in the Evidence field, the study labeled with the first PMID implicates the gene in a biological mechanism/function, and the second one explains how the mechanism/function is associated with PD.

### 8 Supplementary Figures

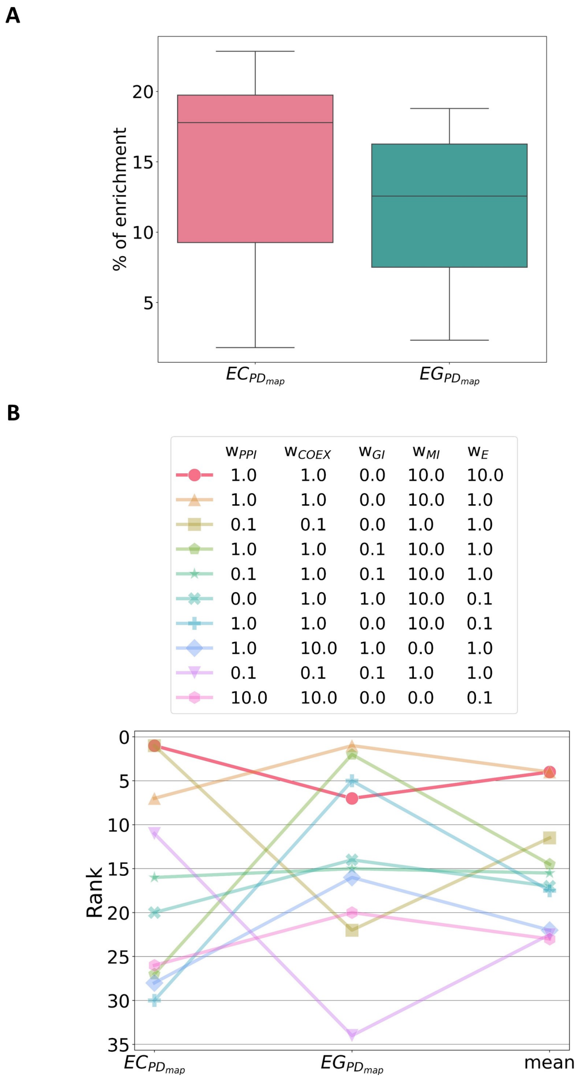


Supplementary Figure 1: **(A) The percentage of enrichment for each enrichment measurement across all combinations of input data. (B) Top 10 combinations of input data.** Each combination of input data is represented by a line where ($w_{PPI}$, $w_{COEX}$, $w_{GI}$, $w_{MI}$, $w_E$) represent the weighting factors by which the corresponding input matrices (PPI, COEX, GI, MI and E) are multiplied. The data combinations are scored by taking the mean of the rank of the gene (EG) and cluster (EC) enrichments in pathways from PD map (PDmap) [3], so that the higher the enrichment is, the lower the rank is.

A

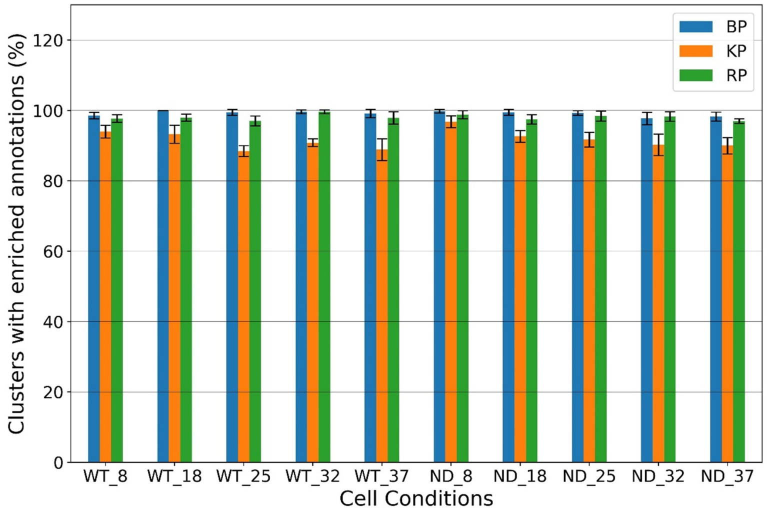


B

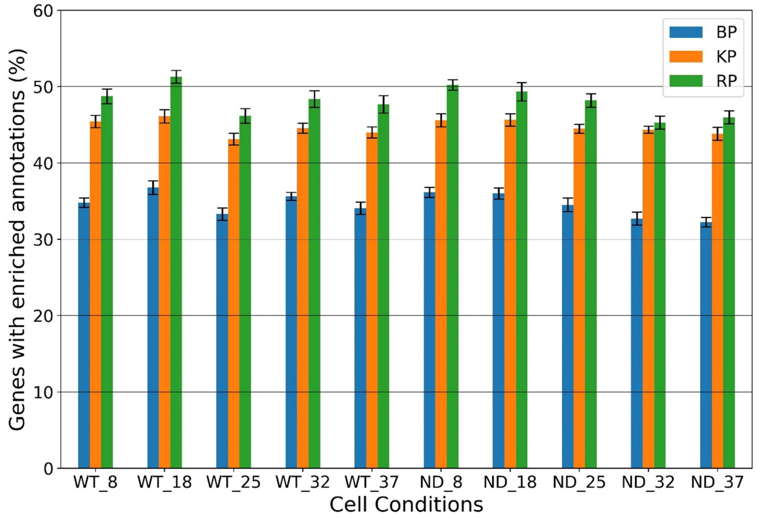


Supplementary Figure 2: **Percentages of (A) enriched clusters and (B) genes with enriched annotations for each cell condition of the best combination of input data, calculated using the matrix $G_1$.** We create clusters of genes for each gene condition and investigate their biological functionality to determine if our integration framework produces biologically relevant gene embeddings. **(A)** For each clustering, the bars show the percentage of clusters with at least one enriched annotation in a cluster, out of all non-empty clusters. **(B)** For each clustering, the bars show the percentage of genes with at least one of their annotations enriched in their clusters, out of all annotated genes. Annotations are KEGG pathways (KP), Reactome pathways (RP), and gene ontology biological processes (GO-BP).

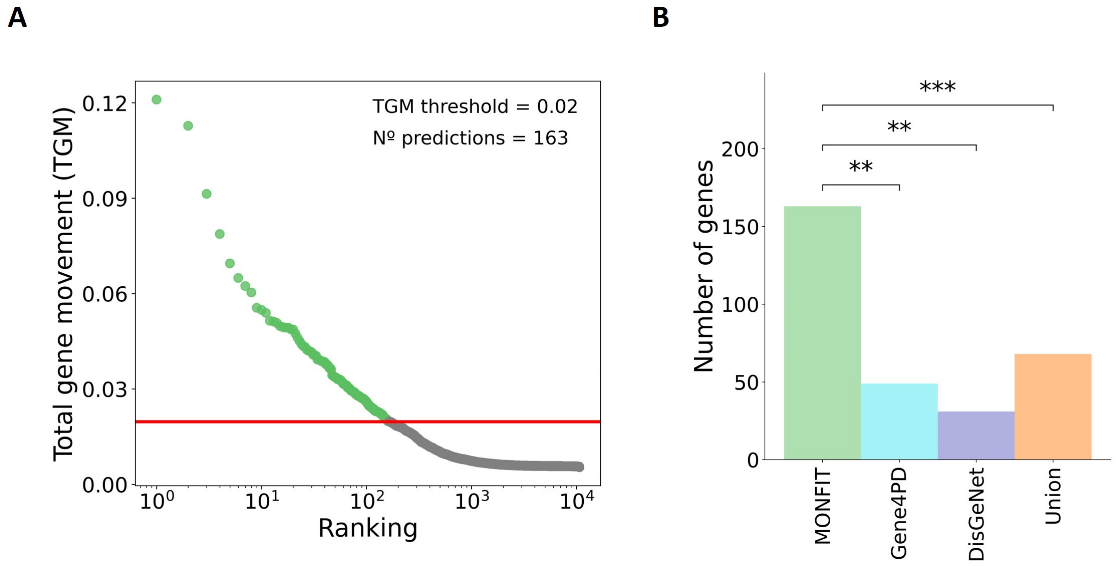


Supplementary Figure 3: **PD gene predictions.** **(A)** Rank-order distribution of the genes according to Total gene movement (TGM). Our PD gene predictions are genes whose TGM values exceed the threshold marked by the vertical red line, resulting in a total of 163 predictions. **(B)** Enrichment of PD gene predictions in known PD genes. We perform the enrichment of PD gene predictions in three sets of known PD genes, obtained from Gene4PD [5], DisGeNet [7] and their union. The bars represent the number of PD gene predictions obtained with MONFIT (green) and the number of those genes that are known PD genes belonging to Gene4PD [5] set, DisGeNet [7] set or their union. The stars above the bars for each comparison indicate the size of the enrichment *p-value* in the corresponding known set of PD genes.

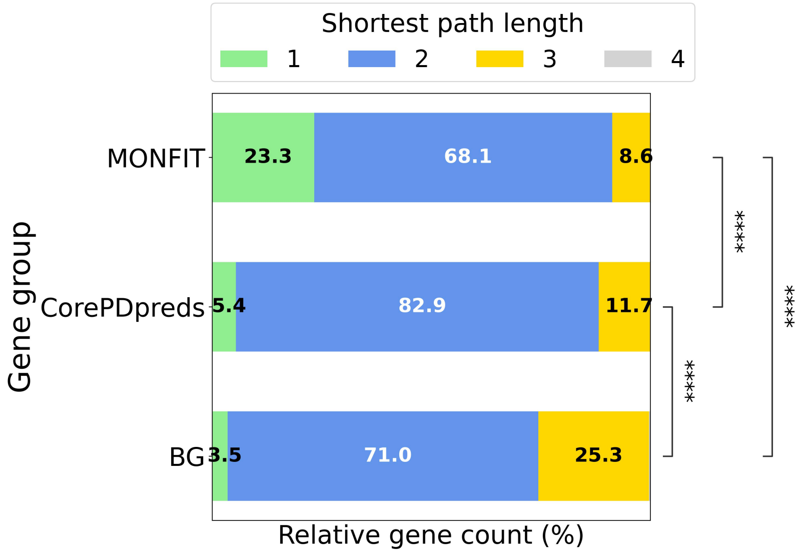


Supplementary Figure 4: **Shortest path lengths to *PINK1* in the PPI networks of MONFIT gene prediction, *Core PD predictions* (CorePDpreds) and background (BG) genes (genes expressed in at least one cell condition, not including the MONFIT gene predictions and *Core PD predictions*).** ***Core PD predictions*** are obtained by applying the 2-step downstream analysis method from Mihajlovic *et al.* [6]. Different patches of one gene set are proportions of genes that are 1, 2, 3 or 4 hops away from *PINK1*. The percentages of genes that are 4 hops away from *PINK1* is $\leq 1\%$, and their labels are not shown in the plot. We compare the shortest path length distributions using a one-sided Mann Whitney-U test. The *p-values* are adjusted for multiple hypothesis testing using the Benjamini-Hochberg procedure [1]. The *p-values* are indicated with stars for each comparison, so that: ******** ($p \leq 0.0001$), ******* ($0.0001 < p \leq 0.001$), ****** ($0.001 < p \leq 0.01$), ***** ($0.01 < p \leq 0.05$), and **ns** ($p > 0.05$, not significant).
